# A Unified Framework for Model-Informed and Agentic RNA Design

**DOI:** 10.1101/2025.06.17.659751

**Authors:** Aidan T. Riley, McKayla Vlasity, Wyatt M. Becicka, Joey Zhuoying Huang, Sicheng Pang, Wilson W. Wong, Mark W. Grinstaff, Alexander A. Green

## Abstract

End-to-end, machine-learning-based design of mRNA molecules offers a powerful means to tailor their properties for specific tasks. mRNA expression level, immunogenicity, tissue specificity, stability, and localization strongly depend on sequence, providing a rich set of properties amenable to optimization. Despite this potential, the components of mRNA are governed by distinct grammatical and functional rules that hinder a unified approach to complete mRNA design. While machine learning and generative AI techniques can excel on individual sequence design tasks, out-of-distribution design, where the biological objective shifts substantially from the original training data, remains difficult. Moreover, there is a disconnect between available sequence generation technologies and the diverse body of biological datasets needed to form and test mechanistic hypotheses. In this work, we describe a simple and powerful alteration to integrated gradients (Design by **I**ntegrated **G**radients or DIGs) that serves as the foundation for several mRNA design tasks and an agentic hypothesis engine, the **St**ructured **R**NA **E**vidence **A**ggregation **M**odule (STREAM), which enables rapid adaptation of this technique to new contexts. Using this framework, we demonstrate complete model-informed mRNA design and reveal the underexplored rules governing the assembly of mRNA components into high-performance transcripts. By linking neural-network-based design to independent datasets, we design complete mRNA sequences in shifted settings, culminating in up to 6-fold increases in intramuscular expression compared to state-of-the-art methods in vivo. Together, DIGs and STREAM enable automated mRNA design in increasingly complex settings.

## INTRODUCTION

Sequence-to-function machine learning techniques show remarkable performance across a range of RNA design tasks^1–10^. From precise engineering of messenger RNA (mRNA) stability^8^ and ribosome load^2^ to the design of novel RNA sensing mechanisms^1,3,4,9^, an increasing degree of control over RNA systems is available through in silico design. Despite these results, a core outstanding challenge for this discipline arises when trying to translate the benefits of model-informed design to experimental settings that were not encountered during training^11^. In particular, approaches that design mRNA 5’ untranslated regions (UTRs), coding sequences (CDS), or 3’ UTRs perform extremely well within their original screening contexts, but struggle when generalizing to new cargos, cell lines, or when they are ported from *in vitro* to *in vivo* models^4,11^. This presents a challenge for therapeutic mRNA design, as screening techniques surveying the span of cell lines, reporters, and animal models necessary to achieve generalization along these axes are unlikely to be available in the near future. In contrast, DNA sequencing technologies have led to the accumulation of a vast body of transcriptomic data profiling countless biological conditions of therapeutic or diagnostic interest^12–14^; however, there are no available techniques for efficiently integrating these datasets with existing sequence-function design tools. Addressing this compatibility gap to mobilize heterogeneous sources of data will generalize mRNA design and unlock unprecedented improvements in mRNA expression, specificity, and immunogenicity in various therapeutic contexts.

This incompatibility is driven by a misalignment between machine-learning-based design techniques and available data. Predictive models can typically be applied and fine-tuned in a straightforward and general manner^4^, but their accuracy sharply reduces with limited data or large changes in experimental context. Generative design approaches built around predictive oracles face an additional, and more challenging, set of constraints. Foundational large language models (LLMs) offer a potential solution to generalize design given their striking degree of success across complex reasoning scenarios^15^; however, directly using LLMs to generate biological constructs end-to-end remains poorly matched to most RNA engineering settings. The power-law scaling governing LLM performance with respect to both available data and computing^16^ is fundamentally misaligned with the task of directly generating biological constructs. Suitable datasets, if available, are often sparse, while new synthetic biology tools and foundation model developments are continuously emerging. Unsupervised language model pretraining can take up to several months on advanced compute clusters^17^, yet fully trained models still require further fine-tuning to align outputs with experimental objectives^18^. Alternative generative techniques such as generative adversarial networks, variational autoencoders, and activation maximization offer a more direct ability to design with respect to a predictive oracle but are intrinsically constrained by the degree of generalization defined by the predictor^4,7,9,19^. Furthermore, these techniques introduce a differentiability requirement throughout their implementation, and, thus, adapting generative model outputs to new grammatical constraints or to extend their design to efficiently account for multiple sources of predictive information is extremely challenging (Fig. 1).

**Figure 1.**
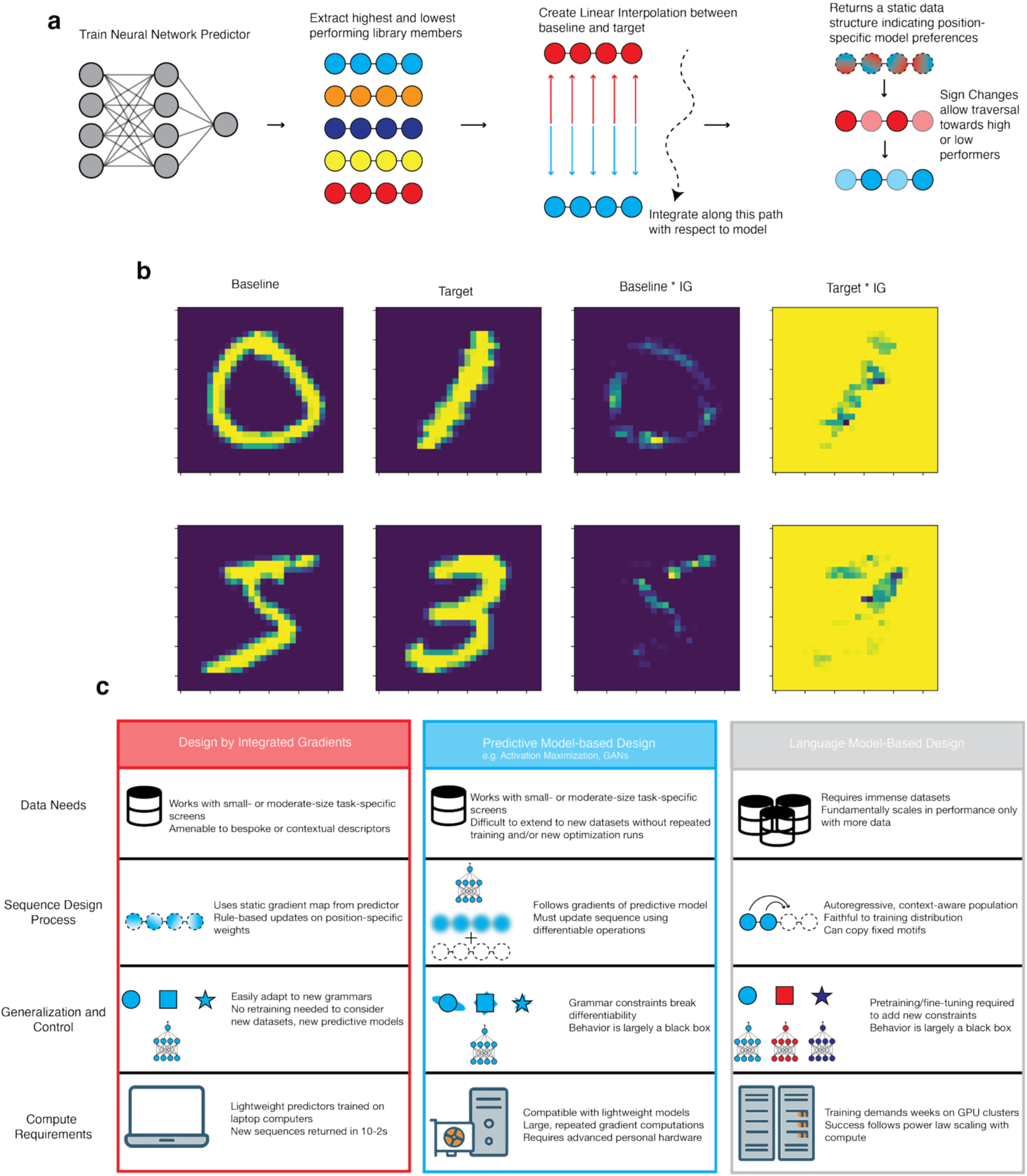
Design by Integrated Gradients. **a**, DIGs utilizes a trained neural network predictor to create a static data structure encoding model preferences. A linear interpolation between the highest and lowest examples within the training dataset is constructed, and the model gradients are integrated along this path. The result of this process is an array in which the sign of individual positions indicates preference/aversion towards the model output. **b,** Application of the DIGs technique to the MNIST dataset of handwritten digits. Element-wise multiplication of the DIGs array by the input reveals specifically which pixels were involved in the class assignment and could be utilized as a template for designing new examples of that class. **c,** Depicted are three different approaches for designing RNA sequences. Design by Integrated Gradients (DIGs, this work) uses a frozen preference array to design sequences, allowing non-differentiable operations to guide nucleotide selection and requiring minimal computing and data resources. Predictive model-based design tools, such as activation maximization or generative adversarial networks (GANs), update sequences iteratively based on the gradients of the model, making it challenging to adapt to outputs to new grammar constraints or new biological contexts. Foundation Model approaches use an autoregressive generation technique to create plausible examples. While generalizable, steering model outputs towards new contexts typically requires fine-tuning and an abundance of data, and outputs risk reproducing encountered training data.

When designing new biological molecules using deep learning methods, each sequence must be represented in a format compatible with gradient-based updates. Discrete symbols (such as A,G,C, and U for mRNA) are therefore mapped to different numerical representations (e.g. one-hot-encoding) that are passed into a model and updated based on the resulting gradient or loss function. Critically, simple edits of bases therefore represent a discontinuous operation in numerical space with no well-defined derivative through which a model can pass information. Several techniques deploy continuously-valued approximations of sequence inputs that can be differentiably updated^2,9,10^. However, strict nucleotide-level control becomes difficult to enforce under these approximation regimes. Accurate CDS translation, targeted structural profiles, and the exclusion or insertion of specific motifs are naturally expressed as discrete, rule-like constraints. In practice, enforcing such grammars while preserving gradient flow typically forces strategies that either (i) encode the rules implicitly into a large generative model during training^1,3,4,6,17,18^, or (ii) rely on soft penalties and post hoc filtering^2,9,10^. The first approach couples biological validity to the model’s learned distribution, narrowing the effective design space, while the second decouples constraint adherence from optimization altogether. Projecting a relaxed, continuous sequence representation manipulated by a predictive model back into a valid discrete nucleotide chain can undo gradient-informed improvements or reintroduce prohibited patterns^4^. Finally, when multiple predictors or loss terms provide competing gradients, suggested updates can interfere, producing unstable tradeoffs or convergence to solutions that satisfy none of the objectives well^4^.

While these challenges of generalization and differentiability affect each portion of the mRNA individually, they are further complicated by the interdependencies of each mRNA component when assembled into a complete transcript. Previous studies outlined the synergistic relationship between naturally occurring UTR elements^20^, while combination screening techniques have revealed the large dynamic range available through UTR selection^21^. De novo, rational design under this combinatorial space of considerations is however severely underexplored due to the lack of machine-guided tools that efficiently design each portion of the mRNA with respect to arbitrary functions.

Unlike generative techniques, various model interpretation methods can produce a fixed map of what types of inputs a model ‘prefers’. While these tools provide a rich understanding of how a black box model constructs its decisions, to the best of our knowledge, these encodings have yet to be explored in the context of generating new outputs. In this work, we investigate a subtle yet enabling modification to integrated gradients^22^ that creates an interpolation between the highest-and lowest-performing members in a dataset used to train our previously published predictive RNA models^4^ (Fig. 2a). This technique returns an array encoding the model’s preferences for specific nucleotides, which when traversed in linear time using different algorithmic techniques affords new sequences. By removing the need for differentiable update steps during optimization, we precisely enforce nucleotide– and grammar-level constraints while still producing sequences with extremely high predicted scores. This degree of control translates the benefits of predictor-supervised design to unseen reporter genes, regulatory mechanisms, and cell lines. Using this toolset to redesign each portion of an mRNA, we apply the concept of coherence to the integration of functional components into a complete mRNA, identifying elements that constructively or destructively affect transcript function. Here, coherence refers to a combination of 5′ UTR, CDS, and 3′ UTR which exhibit a combined regulatory effect reinforcing a shared functional outcome. We further demonstrate that use of these tools and principles increases translational capacity, extends mRNA stability, and improves the dynamic range of RNA regulatory mechanisms in vivo, offering a promising avenue to the challenge of generalizing machine learning-based mRNA design. Finally, we introduce and benchmark an agentic workflow that integrates LLMs as a hypothesis generation layer, rationally separating the reasoning capabilities of generalist foundation models from the responsibility of designing specific nucleotide sequences. In this way, we preserve the interpretable and efficient algorithmic capabilities of our technique and reduce the barrier to entry to efficiently deploy the principles of this work in new experimental contexts. We make the user interface for our design tools and a chat interface for our agentic workflow available at https://rnadesign.app

**Figure 2.**
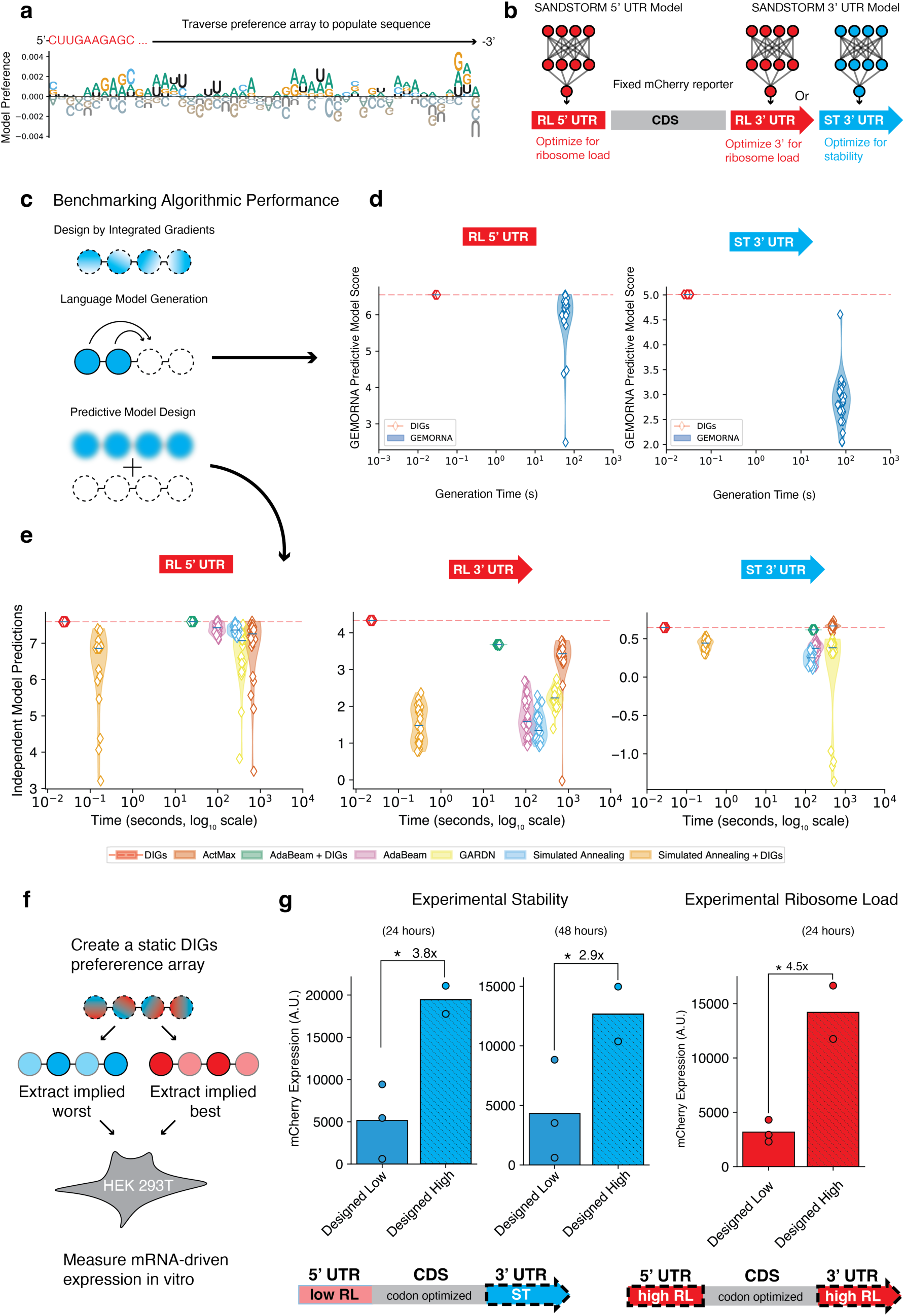
| DIGs UTRs enhance mRNA ribosome load (RL) and stability. **a**, DIGs preference arrays can be utilized as a template for designing new UTR sequences with targeted experimental attributes. **b,** We designed 5’ and 3’ UTRs using pre-trained SANDSTORM models and DIGs to tune the stability and RL of a reporter transcript. **c,** DIGs, large language models, and predictive-model-guided design techniques were each utilized to design new, high-performance UTR sequences. **d,** The inference time (not including pretraining) for language models (GEMORNA^18^) and DIGs are compared on the task of designing high RL 5’ UTR sequences and high stability 3’ UTR sequences. The published predictive models from Zhang et al^18^ were utilized to score outputs. **e,** DIGs runtime is compared to available methods which query a predictive model to return high performance designs. The y axis denotes the predicted performance of individual output sequences as assessed by 10 independent SANDSTORM models that were not directly queried during design (within the range of the normalized values from the original experiment) while the x axis denotes the average runtime of 25 unique designs (averaged over three repetitions). **f,** The dynamic range afforded through DIGs-based design of UTR components was explored. **g,** Engineering 3’ UTR stability through DIGs offers improved expression at 24 and 48 hours for an mCherry reporter protein. DIGs RL UTRs show enhanced expression of an mCherry reporter protein. Data points indicate the mean for an individual 5’ and 3’ UTR combination design across three biological replicates, while bars indicate the mean for the design group. Comparisons denoted * indicate p value < 0.05 comparing each design across groups using independent t-tests.

## RESULTS

### Design by Integrated Gradients (DIGs)

Integrated gradients is a widely utilized approach for unpacking a predictive neural network’s decision-making process^22^. This technique involves the identification of a ‘baseline’ input to a model, typically a zero or noise vector, and the creation of a linear interpolation between this baseline and a ‘target’ sample from the dataset. The model’s gradients are accumulated along this interpolated path, returning an explanatory feature attribution that relates the target’s input features to the model’s prediction. While integrated gradients provide domain-specific insights across diverse predictive modeling use cases, to the best of our knowledge, this technique has not been explored as an engine for designing new data examples with desirable attributes. Thus, we conceptualized an approach whereby we deploy integrated gradients to create a feature map between two samples demonstrating opposite and extreme functions (Fig. 1a) that is carried forward for downstream design tasks. As a demonstration, we first implemented this approach on the MNIST dataset of handwritten images popularly used as a machine learning benchmark. By creating an interpolation between two distinct input examples (Fig. 1a), integrated gradients returned a feature map which characterized the differences between each extreme sample, rather than unpacking an individual element. Further decomposition of this feature map into the regions containing strictly positive or negative values creates a template for designing a new input resembling either the target or baseline (Fig. 1b). Pairing these templates with any number of programming techniques or alternative sources of information guides the extremely efficient, model-informed creation of new samples without the need to retain differentiability. We refer to this technique of pairing an integrated gradients array with downstream design algorithms as <u>d</u>esign by integrated <u>g</u>radients (DIGs). DIGs offers an extremely facile and efficient approach for designing each section of an mRNA molecule with respect to nearly any predictive attribute.

### DIGs designs high-performance 5’ and 3’ untranslated sequences

We first deployed the DIGs approach for mRNA design by creating new 5’ and 3’ UTR sequences designed to impact either the ribosome load (RL) or stability of a reporter protein (Fig. 2). We utilized a SANDSTORM predictive neural network to first map the sequence-function relationship between 5’ UTRs and RL, 3’ UTRs and RL, and 3’ UTRs with stability using previously published datasets^2,23^. After model training, we created a linear interpolation between the one-hot-encoded representations of the highest– and lowest-performing members of the dataset, with the feature attribution returned by integrating along this map carried forward for design. This process entails integrating over a number of interpolated points between both extremes within sequence space. To create a new sequence example using this mapping, we conducted a basic traversal in a 5’ to 3’ direction, selecting the most favored nucleotide at each individual position to maximize a desired attribute and, for a sequence to minimize the attribute, the least favored nucleotide at each position (Methods).

We benchmarked DIGs against a recent, state-of-the-art language model for mRNA design (GEMORNA)^18^ as well as several available design tools that directly query predictive models to guide generation (Fig. 2c). Beginning with GEMORNA^18^, this approach utilized a large-scale pretraining methodology to first create realistic UTR elements, followed by fine-tuning to align outputs with experimental objectives. We tasked this model to return 25 high-performance 5’ and 3’ UTR elements, recording the time for inference only (not including pretraining) (Fig. 2d). For the most direct comparison, we computed a DIGs array using the previous predictive models published in the GEMORNA work (Methods), comparing the inference times and resulting scores of each construct. DIGs decreases inference time by several orders of magnitude while returning 5’ UTRs that match or exceed each of the GEMORNA examples (Fig. 2d). In the case of 3’ UTRs, the quality of DIGs outputs greatly exceeds any of those returned by the language modeling framework (Fig. 2d). In both contexts, the presented improvements in inference time are further compounded by removing the need to first conduct language model pretraining steps when deploying DIGs.

We next evaluated DIGs in comparison to several tools which query predictive models directly to guide generation, including activation maximization^19^, simulated annealing, and our previous work, GARDN^4^ (a generative adversarial network). We additionally evaluated two variations of DIGs that design sequences using different traversal of the preference array (Methods). We similarly tasked each technique with designing 25 individual sequences, reporting the average design time for the set over three repetitions (Fig. 2e).

Analogous to gradient-descent-based training, sequence optimization techniques that query predictive models exhibit overfitting^4^, which leads to cases where the final predicted value of the model utilized during design greatly exceeds that of an identical, independently trained model. This is particularly challenging for activation maximization and generative adversarial network strategies as these techniques are constrained to following the gradient of a single model. In contrast, the static preference arrays returned by DIGs are be averaged across multiple trained predictors, thereby reducing the effects of overfitting while retaining the benefits of gradient-informed optimization. When evaluating the quality of each algorithm’s sequences, we reported the outputs of each technique as the average predicted score across 10 independent SANDSTORM models that were not consulted during design. DIGs designs exhibit the highest predicted performance as judged by a library of independent critic models (not utilized in any design algorithm) on both 5’ and 3’ RL design tasks, while both DIGs and activation maximization return indistinguishable designs for 3’ stability (Fig. 2e). Importantly, DIGs reduces the runtime required for individual sequence design tasks by several orders of magnitude compared to available techniques, setting a new standard for high quality and efficient generation.

To explore the effect of the DIGs approach to tangibly impact the stability of a transcript, we first analyzed the sequence and structural compositions of the resulting designs. In line with a large body of previous literature, the DIGs UTR designed to exhibit minimal stability is nearly completely unpaired in structure while exhibiting a large degree of AU-rich elements known to hinder stability^24,25^ (Supplementary Fig. 1). In contrast, the maximal stability DIGs UTR exhibits a highly folded structural profile and a GC-rich sequence composition which confers degradation resistance^21,26^ (Supplementary Fig. 1). Adhering to known determinants of function along both sequence and structural axes highlights the ability of a simple DIGs traversal to account for features learned by the dual-input SANDSTORM predictive model^4^, an important consideration for designing realistic sequences. We next inserted the DIGs 3’ UTRs into a codon-optimized mCherry expression vector harboring three variations of a 5’ UTR (predicted to have moderate RL). The vector provider (Twist Biosciences) designed both the mCherry coding sequence and the 5’ UTRs. We measured fluorescence generated by translation of the constructs in HEK 293T cells at 24 and 48 hours to assess the impact of stability on total protein expression (Fig. 2f,g). In line with our objectives, incorporation of the DIGs maximal stability 3’ UTR increases expression by 3.8-fold at 24 hours compared to the minimal stability variant. This improvement persists at 48 hours, conferring a sustained 2.9-fold improvement. These results highlight that even a simple computational design approach built around a model-informed encoding provides impactful improvements to mRNA performance.

Several studies report improved protein expression through model-based design of 5’ UTR segments with respect to ribosome load, including our previous work with GARDN and SANDSTORM^4^. To the best of our knowledge, however, the synergistic effects of combining a 5’ and 3’ element both engineered to exhibit maximal RL are unexplored. Thus, we utilized DIGs to design a high RL 5’ and 3’ element using independent experimental datasets of ribosome profiling (Fig. 2g). We again characterized the sequence and structural attributes of these elements, recreating our previous findings of a preference for A-rich sequences with minimal secondary structure incorporated into the 5’ UTR to maximize ribosome load^4^ (Supplementary Fig. 1). By evading differentiability requirements, direct control over nucleotide content is possible in DIGs-designed constructs, allowing consideration of domain-specific grammar during design. This property is advantageous in the creation of high RL 5’ UTR elements, where upstream start codons (uAUG) can potentially hinder translation^27^. We experimentally assessed several DIGs 5’ UTRs with similar degrees of predicted RL that varied in the number of inserted uAUG elements, revealing that multiple uAUG indeed silences translation (Supplementary Fig. 2). Directly excluding inhibitory motifs such as uAUGs while maintaining gradient-informed optimization is a critical capability difficult to guarantee with other techniques that must maintain differentiability. This permits interpretable, grammar-aware sequence design without sacrificing experimental performance.

Despite the availability of suitable datasets, model-based design to reengineer the 3’ UTR for maximal RL is underexplored, leaving the sequence attributes tied to this attribute unclear. Interestingly, we found that our DIGs maximal RL 3’ UTRs displayed a strong preference for U-rich elements and yielded a highly paired secondary structure between the 5’ and 3’ segments corresponding to maximal RL (Supplementary Fig. 1). To assess the degree to which synergistic engineering of RL may impact resulting expression, we evaluated a set of mCherry sequences harboring a DIGs 3’ UTR designed to exhibit minimal RL paired with variations to a standard vendor-designed 5’ UTR (Twist, predicted to have moderate RL) and a second set containing a 5’ UTR and 3’ UTR both optimized by DIGs for maximal RL. The RL designs provide a 4.6-fold improvement in mCherry expression in HEK 293T cells at 24 hours (Fig. 2g), demonstrating the benefit afforded through selection of UTR elements engineered along the same optimization axis. Interestingly, the assembly of a DIGs-stability element with a DIGs-RL element did not demonstrate synergistic effects, but antagonistic decreases in expression. This finding exemplifies one of the limitations of model-based mRNA component design when attempting to generalize to out-of-distribution reporter contexts, encouraging us to investigate into how different design axes contribute to overall protein expression.

### Multi-functional DIGs UTRs enable out-of-distribution performance

The lack of available design tools that integrate diverse sources of information currently restricts the creation of high performance 3’ UTR sequences and is a major bottleneck for generalizing mRNA design (Fig. 3a). 3‘ UTR sequences profoundly alter translation, stability, subcellular localization, and more through an ensemble of RNA binding protein elements, microRNA (miRNA) interactions, and sequence modifications. However, the datasets profiling these interactions are largely siloed. Designing a maximally optimized sequence for a single objective likely comes at the expense of others without explicit accounting such secondary effects. While an immense multi-domain dataset and sufficiently parameterized foundation model may be able to incorporate these diverse sources of information in an unsupervised, ‘top-down’ manner, DIGs offers the direct ability to design with respect to both differentiable models and various sources of domain-specific information in a ‘bottom-up’ approach (Fig. 3a).

**Figure 3.**
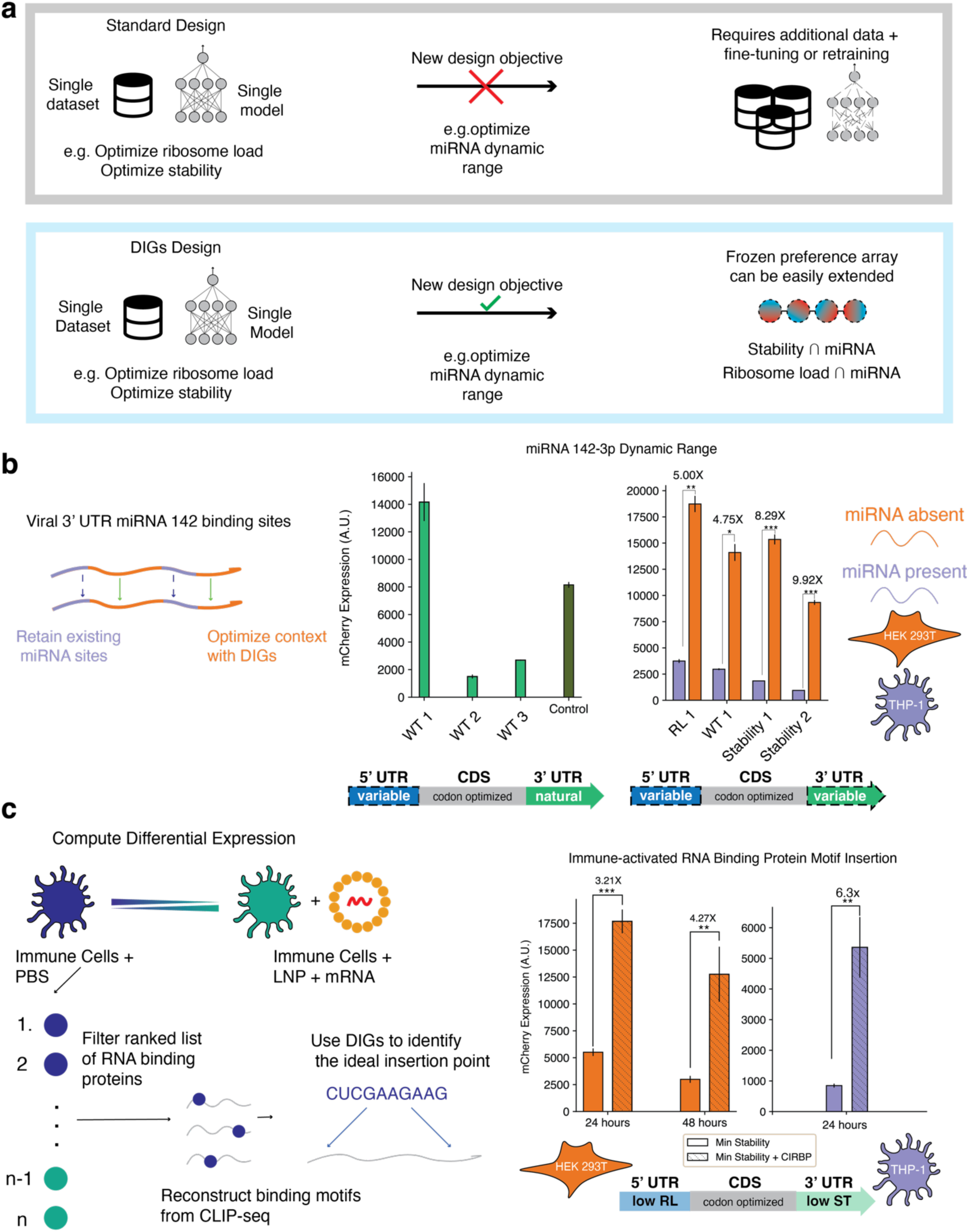
| DIGs incorporates diverse data sources to enable out-of-distribution UTR design. **a**, Extending generative design of mRNA molecules to new functions not directly profiled in high-throughput data is challenging with available tools. The frozen preference arrays utilized by DIGs can be supplemented with deterministic motif inclusion criteria to enable multi-functional UTR designs in the absence of direct screening data. **b,** A naturally occurring viral 3’ UTR was redesigned with respect to either ribosome load or stability while retaining four miRNA-142-3p binding sties. The DIGs-redesigned UTRs improve the dynamic range of miRNA-enabled cell type specific expression between HEK 293T and THP-1 cells (* p<10^−5^, ** p<10^−6^, ***p<10^−7^). **c,** A single-cell RNA sequencing dataset was mined for upregulated RNA binding proteins upon exposure to mRNA components. CIRBP was identified through this process as highly upregulated and a CLIP-seq dataset describing binding preferences was utilized to create a probabilistic reconstruction of the binding motif. The DIGs stability array was utilized to identify the optimal insertion point for two CIRBP binding motifs. The DIGs-guided insertion of these binding elements into our minimum stability UTR element demonstrated a 3.21-fold improvement in expression at 24 hours following transfection and sustained a 4.27-fold improvement at 48 hours post-transfection in HEK 293T cells (*** p<10^−4^, **p<10^−2^). Bars represent the mean expression measurement of an individual design averaged across three biological replicates. Error bars indicate the standard deviation.

As a demonstration of this potential, we explored the integration of the machine-learning-guided DIGs framework with the smallest scale of experimental information available—a functional investigation of a single UTR element^28^. We redesigned a viral 3’ UTR sequence to maximize RL or stability while retaining the endogenous regulatory elements within this sequence (Fig. 3b, left). This UTR, which contains a manually identified quartet of miRNA-142-3p binding sites, confers cellular tropism to this viral strain, silencing expression within certain immune cells^28,29^. Insertion of this UTR into an exogenous RNA vector is expected to confer a similar tropism effect. However, screening the naturally occurring sequence with three variations of a 5’ UTR supplied by a commercial vendor reveals that directly incorporating these components into new vector contexts drastically reduces total gene expression in the absence of the miRNA, limiting their potential in practical settings (Fig. 3b, center). We therefore traversed the DIGs stability and RL arrays as described earlier while bypassing the nucleotide indices corresponding to the miRNA binding sites (Fig. 3a), thus leaving them fully intact while optimizing expression in the absence of the miRNA signal. We reasoned that the resulting mRNA constructs will improve gene expression through either increasing mRNA stability or RL in cells that lack miRNA-142-3p, while remaining silenced in immune cell lines. These designs notably represent a significant departure from the original distribution of sequences utilized to train the predictive models; the original viral UTR is more than double the length of the training samples and previous DIGs designs. In line with our design objectives, the redesigned DIGs UTRs increase the selectivity of an mCherry mRNA vector characterized in HEK 293T (miRNA-142-3p –) and THP-1 (miRNA-142-3p +) compared to the endogenous sequence (Fig. 3b). The redesign constructs improve cell-type selectivity to 5-fold, 8.29-fold, and 9.92-fold, respectively, compared to the 4.75-fold of the highest performing original variant (Fig. 3b, left). These results highlight that DIGs provides an avenue for incorporating bespoke experimental results with the benefits of model-based design without introducing complex fine-tuning techniques.

To explore how descriptive omics datasets enable the generalizability of 3’ UTR design using DIGs, we first explored a single-cell RNA sequencing dataset profiling murine transcriptomic responses to mRNA vaccination^30^. We hypothesized that, within this dataset, upregulated RNA binding proteins could be identified that provide a stabilizing or enhancing effect to endogenous transcripts in this specific context. We further hypothesized that incorporating such a binding motif into an exogenous vector will confer a similar protective effect. In line with this thinking, we identified Cold-Inducible RNA Binding Protein (CIRBP) as highly differentially expressed within the various immune cells of the murine spleen following mRNA vaccination (Fig. 3c, left). CIRBP enhances the stability and expression of different transcripts in various stress response and circadian rhythm pathways^31–33^, with putative binding sites identified through CLIP-seq screening techniques^34^. We carried the probabilistic reconstructions of the CIRBP binding motif (Methods) forward for DIGs-based design, hypothesizing that the optimal placement of these elements within a 3’ UTR will further enhance improvements in vector stability in a context-specific manner.

To incorporate these elements into a DIGs design, we employed an iterative greedy algorithm to locate the ideal insertion point of each regulatory motif. This entailed scanning each regulatory element against the DIGs stability or RL array from a 5’ to 3’ direction and calculating the implied score of inserting the motif at each position, eventually choosing the highest score for final placement. Recent language modeling techniques have deployed similar approaches to rationally insert known motifs into model-designed UTR elements, demonstrating no improvement in performance^18^, leading to the conclusion that 3’ UTRs confer a secondary role to total mRNA performance. As shown in Figure 3d, DIGs-guided insertion of the CIRBP binding sites recovers the translational capacity of the minimal stability element by 3.21-fold at 24 hours and 4.27-fold at 48 hours. This improvement is enhanced even further to 6.3-fold when evaluating the same pair in an immune-competent cell line, THP-1, confirming that this approach bridges diverse datasets and probabilistic sequence analysis tools with the benefits of model-informed design (Fig. 3c, right. Taken together, these findings demonstrate how DIGs bridges sequence design informed by *in vitro* screening into more complex, out-of-distribution contexts necessary for RNA therapeutics.

### Combinatorial screening of DIGs UTRs reveals functional rules for selecting UTR pairs

While these results are useful for translating individual UTR elements to out-of-distribution tasks, mRNA performance is determined not only by individual RNA components (e.g. UTRs and CDS) but also by how these components interact with one another in the mRNA and the larger mRNA interactome. However, the rules that govern mRNA component assembly are not well understood. A substantial body of previous literature concluded that engineering mRNA molecules to exhibit maximal stability is the ideal approach when optimizing for overall protein expression^5,8,18,19,35–42^. Specifically, maximal RL leads to poor expression through translation-dependent degradation^35^, while moderate or minimal RL of a highly stable mRNA mitigates this effect^5^. These highly stable sequences predominantly recruit monosomes, evading the translation-dependent degradation pathway while prolonging the lifetime of the transcript and increasing cumulative expression over time^18^. In contrast, high RL leads to a bursting expression pattern at early time points, ultimately leading to less total protein over time^18^. This perspective is the basis for many sequences that are actively being explored within the clinic or are already approved for treatment^43^.

Given the lack of tools that effectively design each component of the mRNA simultaneously, we hypothesized that the synergistic effects of extreme RL and stability-based UTRs may be incompletely characterized (Fig. 4a). Understanding this interaction term will provide insight on why the insertion of individually optimized UTR elements into alternative vectors tends to limit performance. We screened a small library of DIGs UTR elements designed to effectively survey the extremes of these attributes with a fixed reporter sequence. This library consists of 3’ UTR elements designed to have maximum RL, maximum stability, minimum RL, or minimum stability, as well as 5’ UTRs designed to exhibit maximum RL and 5’ UTRs predicted to have moderate RL commonly used in Twist Biosciences vectors. Importantly, the sequence-function mappings guiding the creation of each of these components originate from different experiments, functional assays, and laboratories, leaving the principles guiding their rational selection completely open-ended (Fig. 4a). We assessed this library at multiple time points to potentially identify distinct temporal profiles offered through each attribute. The expression profiles of the sequences demonstrate a wide dynamic range at 6, 24, and 48 hours post-transfection, between the highest and lowest performers (Fig. 4b), highlighting the importance of the rational assembly of components to design complete, high-performance mRNAs.

**Figure 4.**
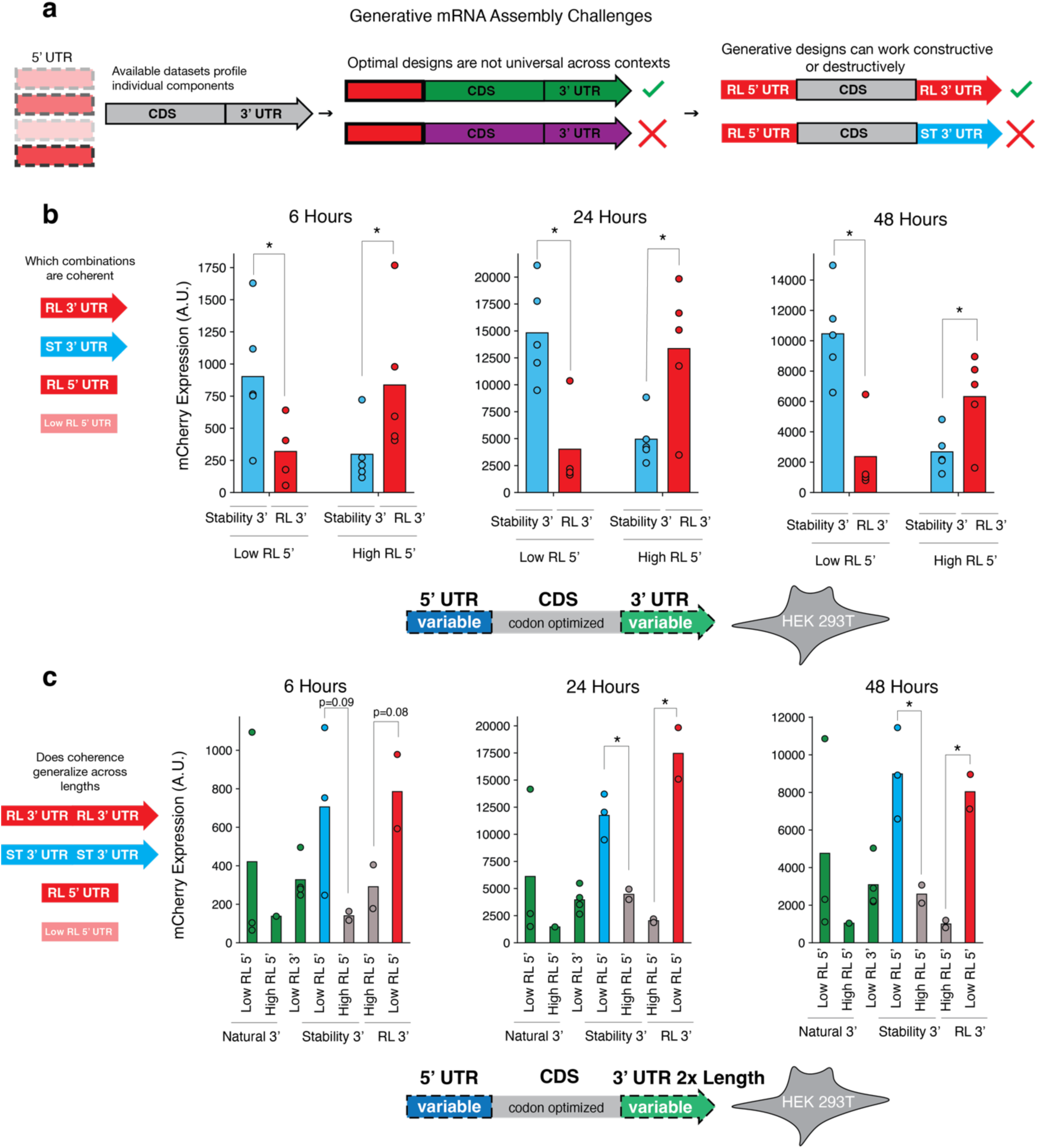
| Combination screening reveals coherent UTR selection criteria. **a**, High-throughput screening is constrained to libraries of individual components. Model-designed sequences can inhibit performance when ported into new reporter constructs or when combining members from differing screening libraries. **b,** A library of engineered DIGs 3’ UTRs was assessed in combination with DIGs and Twist 5’ UTRs in HEK 293T cells. mCherry expression at 6, 24, and 48 hours is reported. Comparisons denoted * indicate p < 0.05 measured using Brunner-Munzel non-parametric statistical test. **c,** The coherent selection criterion remains informative when considering miRNA-142 redesigns which harbor four fixed miRNA binding sites and are double the length of original DIGs designs. Comparisons denoted ‘*’ display p < 0.05 measured using Brunner-Munzel non-parametric statistical test. Comparisons denoted ‘*’ display p < 0.05 measured using independent t-test for each pairwise sample comparison within the groups across three biological replicates. Data points in all plots represent the mean expression for an individual design across three biological replicates. Bars represent the mean measurements from a specific group of designs. Error bars indicate the standard deviation of the design group.

Profiling this library of elements revealed a surprising result – combining a 5’ and 3’ element designed for maximum RL creates a robust expression profile over time rather than demonstrating clear signs of translation-dependent degradation. The same expression improvement is also seen when selecting a 5’ element for moderate RL and a 3’ element for maximum stability. In contrast, the combinations of elements designed to exhibit extreme but mismatched traits (such as high RL 5’ and high stability 3’) nearly universally decrease expression (Fig. 4b). This result may explain the stability-focused findings of previous studies, where the placement of a high RL 5’ UTR within an otherwise unmodified transcript counterintuitively reduced total protein expression^21,35^, while highly structured 5’ UTRs that limit ribosomal binding increased total protein output. For the remainder of this paper, we will refer to combinations of mRNA elements designed to emphasize the same attribute (e.g. RL or stability) using the term coherent selection, while combinations of RNA elements where there is a mismatch in attributes will be termed incoherent selection.

A time-course analysis of various UTR elements at 6, 24, and 48 hours post-transfection reveals an additional surprising finding: a simple coherent selection process effectively optimizes total protein over time, regardless of the axis for design. While we identified a slight shift in expression favoring the stability designs at 48 hours, no significant differences are present between the orthogonal design approaches in terms of fluorescence AUC calculated over all time points, while most combinations of incoherent UTR elements show minimal expression. Individual UTR combinations additionally validate that designs created through stability engineering still demonstrate high early expression, while those designed through RL persist despite translation-dependent degradation effects^35^ (Fig. 4b,c). This benefit of coherent UTR selection additionally held when considering our synthetic miRNA-142 redesign constructs (Fig. 4c), which fall greatly outside the length distributions of the original datasets used for model training, indicating the generalizability of this principle. To further investigate the ubiquity of UTR coherence, we screened a subset of the coherent and incoherent design UTR pairs in THP-1 cells, an immortalized human macrophage line (Supplementary Fig. 3). Interestingly, while the coherent pairs still outperformed their incoherent counterparts along both axes, a strong preference for the stability-based vectors exits alongside a relative reduction in the RL-based designs. This finding suggests that individual cell types may exhibit preferences for tunable design objectives; a detailed understanding of this principle will enhance model-based mRNA design in out-of-distribution experimental settings and potentially serve as a lever for engineering cell type specific expression.

### Transcriptomic analyses challenge the assumptions of sequence design paradigms

Building off our initial UTR findings, we next uncovered if various transcriptomic datasets could be mined for informative signals to enhance model-based mRNA design in diverse settings. A robust body of literature details the benefit of ubiquitously designing coding sequences and 3’ UTRs to exhibit the most extensive secondary structure possible^8,18,21^ while incorporating the most abundant codons possible. A number of *in vitro* experiments, which demonstrate maximally folded mRNA molecules are more stable against hydrolysis and exhibit more prolonged half lives in immortalized cell lines,^8,41^ provides the motivation for this design objective. This paradigm does not, however, consider the dynamic impact of innate immune sensing present within *in vivo* models, tissue-specific differences in codon abundance, or the potential synergistic effects of coherent mRNA component selection on overall expression. Given the dynamic variance of these attributes across the settings in which mRNA could be useful, refining our understanding of how these factors should be incorporated into design remains a critical outstanding challenge.

While increased immune stimulation may be advantageous for vaccine development, there are many therapeutic applications for mRNA in which immune activation must be avoided^42^. Furthermore, the interaction between total expression of an RNA payload and dsRNA-activated immune degradation is understudied in the context of design. We conducted an in-depth analysis of transcriptomic changes following exposures to different types of RNA to assess whether a global preference for different structural profiles or codon usage contributes to differential expression (Supplementary Fig. 5). Beginning with a dataset mapping the time-course transcriptomic response of human macrophages to Chikungunya infection^44^ (an RNA alphavirus^45^ adapted for RNA-based therapeutics), we found that the length-normalized minimum free energy (MFE) of up-regulated transcripts is significantly greater than that of down-regulated transcripts across both 8 and 24 hours (Supplementary Fig. 5a). We continued this analysis using a dataset characterizing the tissue-specific transcriptomic response to Chikungunya infection in both macaque and mouse models^46^. In these *in vivo* settings, a similar association of minimally structured upregulated genes is seen in both the lungs and spleens of these animals, while the liver demonstrates the opposite trend in both animals (Supplementary Fig. 5c). Of the tissues analyzed, the heart is the only example that demonstrates a preference for less-structured transcripts in macaques, but no significant difference in mice. These findings suggest a more nuanced approach to the design of coding sequences for *in vivo* applications than pure secondary structure maximization.

To isolate this effect from potential impacts of viral proteins on immune response, we then characterized the response to an mRNA vaccine enclosed within a lipid nanoparticle (LNP) at single-cell resolution in mice^30^. We computed differential expression of cells within the excised muscle of mice exposed to a COVID-19 vaccine mRNA enclosed within an LNP and those exposed to an empty LNP, revealing a striking increase in length-normalized MFE for upregulated genes across nearly every cell type (Supplementary Fig. 5b). Interestingly, the structural differences between up– and down-regulated genes do not vary as a function of spike mRNA count, indicating a global effect at the site of vaccination. Furthermore, when isolating the upregulated genes of specific cells, a clear difference exists between immune and muscle cells, indicating that the optimal composition of codons, which dictates much of the mRNA MFE, may differ depending on the cell type and condition of interest (Supplementary Fig. 5d). Understanding the exact role of mRNA codon usage and secondary structure in a context-specific manner will greatly enhance our ability to execute rational mRNA design in previously unobserved settings, but a detailed explanation of these contributing factors remains unobservable with existing tools which explicitly prioritize increasing structure while minimizing RL.

### DIGs CDS designs uncover the contributions of diverse factors on mRNA performance

While these results provide a motivation to explore the relationship between secondary structure, ribosome load, and context-specific forward design of coding sequences, available CDS design approaches are typically formulated to maximize secondary structure and species-wide codon preferences (Fig. 5a). Furthermore, adherence to an arbitrary amino acid sequence poses a challenge for gradient-based design techniques that are further limited to varying each of these properties individually (Supplementary Fig. 5). In contrast, DIGs is compatible with arbitrary predictive oracles, accounts for grammar in a flexible manner, can be formatted to parse out the explicit contributions of secondary structure, codon usage, and UTR selection on overall mRNA performance (Fig. 5b,c).

**Figure 5.**
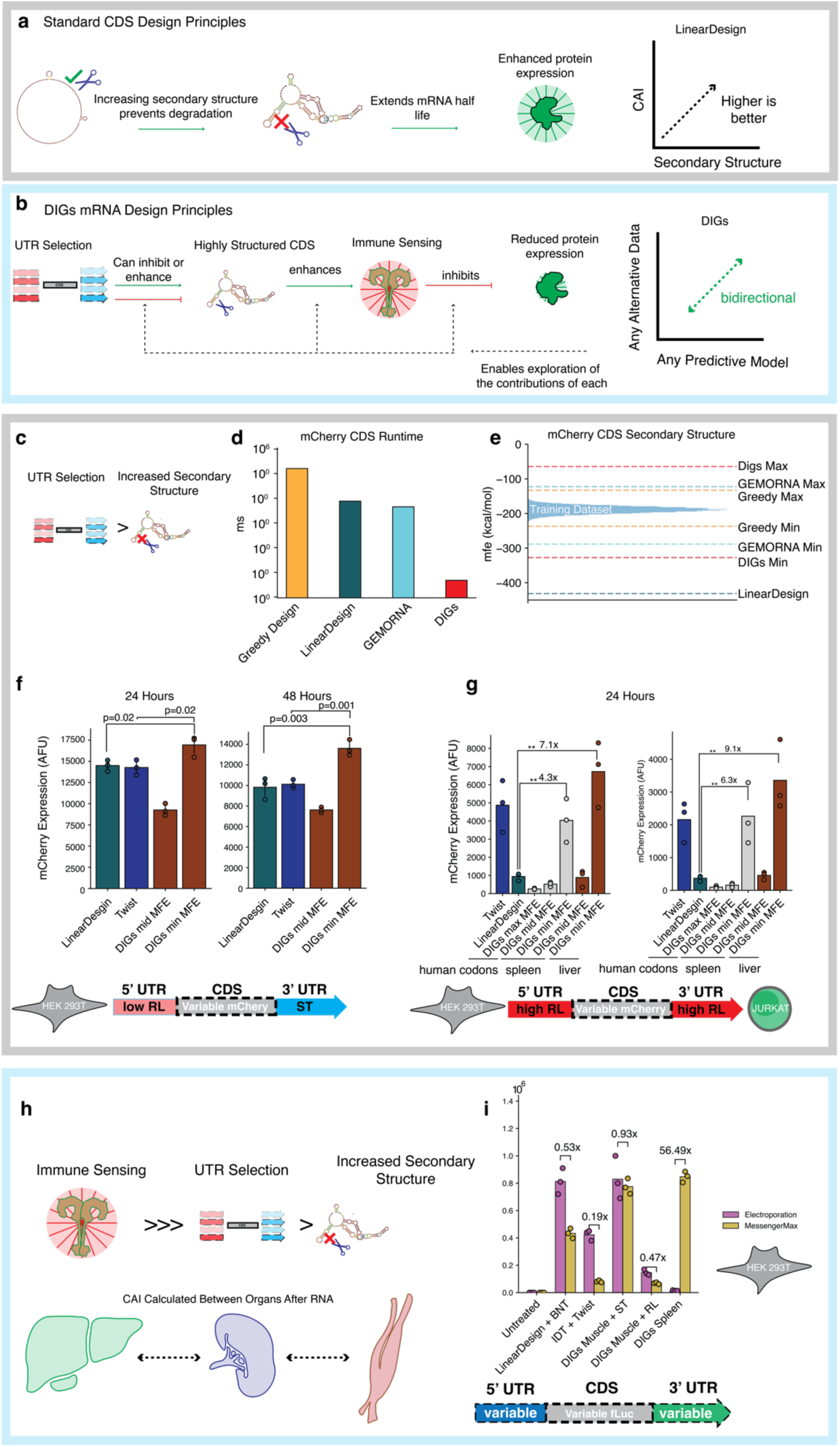
| DIGs-driven CDS optimization reveals mechanisms beyond secondary structure and CAI. **a**, Standard CDS design approaches leverage the relationship between secondary structure and RNA half-life to design optimal candidates, prioritizing high CAI values and increasing degrees of structure. **b,** DIGs is extensible to arbitrary predictive models or non-differentiable suggestions, allowing exploration of a greater range of the space of possible designs. These attributes contribute to a more complete model of which attributes enhance protein expression from mRNA molecules. **c,** Coherent selection of CDS and UTR elements along the same design axis outweighs structural maximization when optimizing total protein output. **d,** DIGs-based design is extremely efficient, enabling several orders of magnitude improvements in design time reductions compared to LinearDesign, GEMORNA (not including pretraining), or an analogous greedy approach. **e,** DIGs forward designs extend beyond the structural attributes of the sequences observed during training, and extend further than GEMORNA or greedy design but not as far into low MFE levels as LinearDesign. **f** DIGs coding sequences designed to exhibit maximal secondary structure while utilizing the codon preferences of the liver following mRNA exposure shows improved expression at 24 and 48 hours compared to either LinearDesign or twist. Bars indicate the mean expression measurement across three biological replicates. **g,** The CDS library was evaluated for expression using the optimal coherent RL pair in both Jurkat cells (electroporation) and HEK293T cells (MessengerMax, methods). Bars denote the mean expression for a single design across three biological replicates while points denote individual replicates. **h,** Immune sensing triggered by endosomal uptake of RNA is a primary consideration for designing therapeutics. **i,** DIGs mRNA constructs designed with contextual data describing cellular response to RNA exposure were profiled with controls using electroporation and MessengerMax. While MessengerMax reduces the efficiency of most designs, the DIGs spleen-specific construct shows a dramatic 56-fold increase in expression through endosomal delivery. Bars indicate the mean luciferase luminescence recordings over three biological replicates, while markers indicate individual replicates.

We investigated this challenge by creating a library of synonymous coding sequences that varied based on their degree of secondary structure and their use of context-specific codon preferences. To create MFE-varying CDS designs using DIGs, we first utilized a suite of SANDSTORM predictors to estimate the minimum free energy (MFE) of various CDS datasets. We created these datasets by sampling hundreds of thousands of examples of the same protein coding sequence in silico (e.g., mCherry, green fluorescent protein), using uniform codon biases and were labelled according to their length-normalized MFE calculated by thermodynamic models (Methods). We trained each predictive model on an individual synthetic protein dataset and queried using integrated gradients between the highest and lowest MFE examples. This array therefore contains globally informed, position-specific information related to the secondary structure of possible CDSs (Supplementary Fig. 6) that traverse in linear time with respect to the target amino acid chain.

Unexpectedly, a simple greedy algorithm selecting the highest-scoring valid codon suggested by this array efficiently generalizes beyond the initial training dataset of 500,000 sequences when tasked with designing either extremely highly or lowly structured sequences (Fig. 5e). DIGs additionally generalizes far beyond an analogous greedy approach that directly selected codons using a thermodynamic model informed over the entire sequence. This emergent property reflects the advantage of utilizing globally informed information encoded within the position-specific array returned by the DIGs approach. When compared to LinearDesign set to ignore codon biases, mCherry sequences returned by a simple greedy selection using DIGs do not reach the same extreme minimum. However, the traversal of the DIGs array could in principle be extended to the more sophisticated lattice parsing approach that empowers LinearDesign.

To emphasize how grammatical constraints stress machine-learning-based sequence design tools, we conducted similar CDS design experiments using activation maximization under the supervision of our trained structural predictor, and unsupervised generation using GEMORNA^18^. In the former, activation maximization returns RNA sequences with meaningfully altered structural profiles but only consist of an average of 23 / 237 correct amino acids; we therefore omit activation maximization from further consideration (Supplementary Fig. 5). In contrast, GEMORNA learns the RNA translational grammar in an unsupervised manner, but the next-token prediction formatting that guides GEMORNA pretraining is fundamentally disconnected from tangible sequence attributes and limits generated diversity. To illustrate this point, we generated 10,000 examples of mCherry CDS elements, 2,000 examples of mCerulean, and 2,000 examples of Nano-luciferase. In each case, the publicly available GEMORNA model returns only 200 unique elements, which could be the product of over-constrained decoding or mode collapse.

The runtime of DIGs additionally outperforms either LinearDesign, greedy design based on thermodynamic models, or GEMORNA by several orders of magnitude while allowing bidirectional optimization with a simple python implementation (Fig. 5d). Furthermore, while we explore secondary-structure-based design of the CDS using DIGs, this approach accounts for CDS-specific grammar while designing with respect to arbitrary differentiable predictive models. This feature will be increasingly useful as DNA screening technology for extended library lengths develops and direct sequence-function readouts of CDS elements become possible.

Unlike other methods that could potentially design a sequence with respect to a predictive neural network, DIGs traversals are easily supplemented with arbitrary, non-differentiable sources of information. We leveraged this feature to simultaneously incorporate context-specific codon usage metrics into our designs while extending the degree of CDS structure significantly beyond encountered training data (Supplementary Fig. 6). This entailed calculating the most upregulated genes in a tissue and condition of interest, collating the codon usage of these genes into a unified codon adaptation index (CAI) that could be queried, and supplying these suggestions in parallel to the structural considerations to the DIGs traversal during runtime (Methods). Our final library therefore contained seven CDS variants of mCherry. Three DIGs designs used the codon preferences of the murine spleen (encompassing AT-rich codon preferences) following chikungunya exposure and demonstrated varied structures of –64.4 kcal/mol, –87.5 kcal/mol, and –215.8 kcal/mol. These CDS elements exhibit minimal secondary structure and represent a class of coding sequence that are unachievable using other available techniques. An additional two DIGs designs utilized the codon preferences of the liver in this same experiment (encompassing GC-rich codon preferences), demonstrating MFEs of –312.7 kcal/mol and –323.5 kcal/mol and thus higher degrees of secondary structure. The final two library members were a standard, codon-optimized mCherry sequence from Twist and a construct returned by LinearDesign with MFE values of – 272.5 kcal/mol and –431.3 kcal/mol respectively.

We assessed the DIGs liver and LinearDesign constructs using the coherent UTR pair consisting of the full-length twist 5’ UTR and stability-based DIGs redesigned 3’ UTR containing miRNA-142-3p binding sites in HEK 293T cells. We hypothesized that this group of elements constitute the most direct extension of previous CDS design approaches, which emphasized moderate-to-low ribosomal loading and stability maximization, while incorporating our UTR selection principles and out-of-distribution specificity offered through miRNA-142-3p binding. The lowest-MFE (min MFE) DIGs design created with context-specific codons exhibits the maximal degree of expression at both 24 and 48 hours, followed by LinearDesign and Twist, and the mid MFE DIGs design (Fig. 5f). Intriguingly, the optimal min MFE DIGs construct outperforms LinearDesign despite being extensively less structured, indicating that context-specific codon selection and optimal UTR combinations supersede structure when optimizing for expression. The increase in expression at 48 hours further indicates that effective degradation resistance can be conferred through proper UTR selection. The performance of this vector highlights how individual experimental findings, context-specific descriptive datasets, and machine-guided design can be unified to create high performance mRNA vectors in the absence of direct high-throughput screening data.

To uncover how the principles of coherent component assembly, CDS secondary structure, and context-specific codon composition govern the performance of mRNA in a more generalized manner, we evaluated each of the CDS elements with the optimal combination of RL UTRs described earlier in both HEK293T and Jurkat Cells (Fig. 5g). Interestingly, each of the minimum MFE designs composed of the splenic and liver codon profiles perform well in both contexts, while the Twist CDS element (standard codon optimization) retains the high performing expression profiled in our earlier coherence screens (Fig. 3.8a,b). As expected, CDS elements designed to be nearly single-stranded using DIGs demonstrate minimal expression (Fig. 5g). while the relative performance of LinearDesign decreases dramatically compared to our earlier demonstrations utilizing the UTRs from the stability axis. In this context, the optimal DIGs CDS elements increase expression up to 9-fold compared to LinearDesign, demonstrating that coherent component assembly indeed applies to the design of both coding sequences and UTRs, and that maximization of CDS secondary structure can be antagonistic to total expression.

mRNA therapeutics typically require cellular uptake through the endosomal pathways^47^, a process which is known to stimulate innate immune activation through pattern recognition receptors and dsRNA sensing mechanisms^48^. The expression-maximizing degree of secondary structure in an mRNA CDS therefore must consider a tradeoff between extending molecular half-life and evading immune activation, yet available tools singularly prioritize an increase in structure. To evaluate which objectives are the most impactful under endosomal uptake contexts, we created a new library of firefly luciferase CDS elements using the CAIs of differing murine tissues following mRNA exposure (liver, spleen, muscle, see methods) and optimal UTR components. We separately transfected the resulting mRNA into HEK 293T cells using electroporation (thereby evading the endosomal compartments) and MessengerMax (leveraging endosomal uptake) (Fig. 5h,i). In this context, LinearDesign + BNT shows a decrease in total protein output of 0.53-fold with MessengerMax transfection compared to electroporation, while IDT CDS + Twist UTRs shows an even greater reduction of 0.19-fold (Fig. 5h). Similarly, the DIGs Muscle + RL constructs demonstrate a 0.47-fold reduction, while the stability constructs exhibit only a 0.93-fold reduction in expression (Fig. 5f). In contrast, the DIGs spleen design shows a 56-fold increase in luciferase activity under MessengerMax transfection conditions, increasing from near baseline in electroporation conditions to the highest total protein output within the group (Fig. 5i). This constitutes a greater than 100-fold improvement in fold-change compared to LinearDesign, despite the substantial difference in their respective degrees of structure (−(-1109 kcal/mol for LinearDesign, -687 kcal/mol for DIGs Spleen), indicating that structure is not the decisive optimization objective to increase total protein expression, and providing a promising route to design optimal therapeutic sequences in more complex biological settings. These factors culminate in a more complete understanding of the objectives governing mRNA performance and will accelerate the transition of model-informed design into increasingly complex biological settings.

### In vivo validation of DIGs constructs

To further validate the generalizability of end-to-end mRNA design using DIG, we created a new set of firefly luciferase benchmark sequences and evaluated them in *in vivo*. In vaccine settings, maximizing expression of an mRNA in both muscle tissue and the draining lymph nodes is essential to achieve robust clinical protection^47^. We therefore created a firefly luciferase (fLuc) CDS using DIGs that jointly maximized secondary structure and the CAI of the most upregulated transcripts in murine muscle tissue following vaccination. We evaluated this CDS with two sets of UTRs. The first consisted of the stability axis 5’ UTR evaluated throughout this work paired with a hybrid 3’ UTR created by stitching together the CIRBP-harboring UTR (hereby referred to as combination) with the coherent stability construct. The second consisted of the RL axis 5’ UTR evaluated throughout this work, with a similar 3’ UTR composed of the stitched combination and RL 3’ UTR. Strikingly, whole-body flux measurements at 24 hours post-intramuscular injection reveal 6-fold and 16-fold improvement in expression for the DIGs Muscle CDS + stability UTR mRNA compared to LinearDesign and IDT mRNAs, respectively (Fig. 6a). This improvement persists across extended time courses, where the DIGs design increases expression 4.3-fold and 6.4-fold at 72 hours post-injection (Fig. 6a). AUC analysis of the expression curves for the optimal DIGs constructs reveals a 3-fold and 10-fold total improvement, respectively compared to the control groups. Interestingly, in muscle tissue, the coherent RL DIGs constructs perform less favorably than the stability UTR pair, indicating a preference for the stability axis in vaccination contexts.

**Figure 6.**
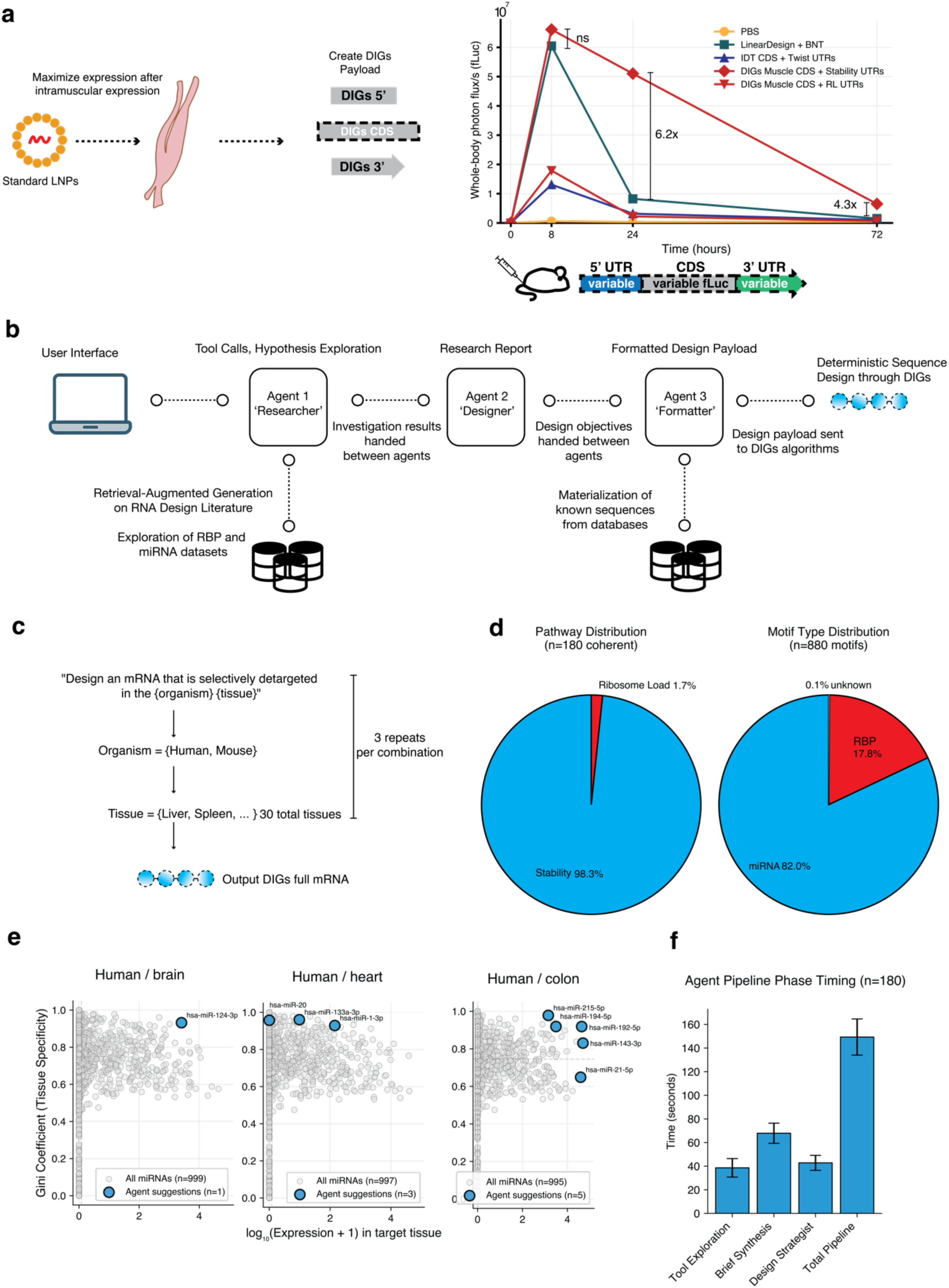
| Context-specific, end-to-end mRNA design using DIGs and STREAM. **a**, A library of nLuc designs were assessed *in vivo* via intramuscular injection with standard LNPs. Whole-body luminescence recordings collected 8– and 24-hours post injection are reported. Data was collected from n=5 mice. Bars denote the mean luminescence values while individual markers indicate recordings from individual animals. **b,** STREAM is a multi-phase agentic workflow that converts user prompts into tailored parameters compatible with deterministic DIGs designs. STREAM is supplemented with miRNA^50^ and RBP databases^51^ as well as RAG using RNA design literature. **c,** A benchmark was conducted across 30 randomized tissues of humans and mice drawn from the miRNA tissue atlas dataset^50^ using a standardized prompt in triplicate. **d,** The suggestions of the STREAM workflow were quantified in terms of the distribution of stability vs. RL UTR selections that were made as well as miRNA and RBP incorporations across each of the 180 distinct design runs. **e,** Three tissues with well-known characteristic miRNAs were analyzed more closely to evaluate the quality of STREAM detargeting motif suggestions. **f,** The average time of complete multi-phase design runs were calculated. Bars denote the mean across n=180 total replicates, while error bars denote the standard deviation.

### Automated RNA design using STREAM

These results spanning CDS and UTR optimization highlight the capability of DIGs to unify rational design principles with model-guided optimization. Still, parsing through available literature and datasets for actionable design signals remains a challenge dependent on deep domain expertise. To generalize these capabilities across contexts, we developed a user interface for the DIGs framework and an agentic workflow powered by generalist large language models to formulate novel RNA design hypotheses. This agentic system, which we refer to as **ST**ructured **R**NA **E**vidence **A**ggregation **M**odule (STREAM) (Fig. 6c) consists of a three-phase process for converting natural language prompts into optimized mRNA molecules. Phase 1 uses retrieval augmented generation^49^ to supplement user queries with a database of tailored RNA design literature (consisting of the totality of references cited here). Following this, STREAM can explore a curated set of data artefacts that guided the creation of each out-of-distribution construct explored in this work. miRNA expression atlases^50^, RNA-binding protein motif datasets^51^, cell-type specific codon profiles, as well as each precomputed DIGs array can be explored, with the reasoning capabilities of the agent guiding the selection of pertinent data to include. Phase 1 terminates with a design report that is returned to the user, highlighting a summary of the exploration and key findings that can be incorporated into the workflow.

This same data report is next handed off to a subsequent agent in phase 2, where these findings are mapped to the abstract principles that guided effective design in this work. This agent is prompted to consider coherent UTR selection, structural tradeoffs, immune sensing, cell-type differences, and effective motifs to include to achieve the user’s design objective, eventually suggesting an overall design approach for each of the 5’ UTR, CDS, and 3’ UTR. These suggestions are finally propagated to phase 3, where each finding is mapped to a specific DIGs parameter. At this stage, deterministic evaluation of suggested motifs, DIGs arrays, and other findings is conducted to ensure auditability in the resultant hypothesis. The formatted outputs of the workflow are finally passed to the DIGs framework for design execution. By deploying STREAM in this manner, open-ended reasoning related to a biological objective is decoupled from deterministic and efficient sequence generation provided by DIGs.

As agentic systems can suffer from hallucinations, we conducted a suite of benchmarks to evaluate the capabilities of STREAM. A standardized prompt was deployed in triplicate across 30 randomly selected tissues or cell lines available in the miRNA tissue atlas for both humans and mice (Fig. 6d). The final DIGs design parameters suggested by STREAM were evaluated for their accuracy along a number of different axes. First, the suggestions of coherent/incoherent UTR combinations were profiled (Fig. 6e), revealing that STREAM adheres strictly to the principles of coherence when suggesting design objectives for both UTR components across all instances. Interestingly, retrieval-augmented generation using RNA design literature noticeably biases STREAM towards the stability axis; coherent ribosome load UTRs are only suggested in 3/180 (Fig. 6e) instances while the remaining suggestions favor the stability axis, highlighting the previous challenges of exporting high ribosome load constructs into out-of-distribution settings.

Following this, the motifs selected by STREAM to confer a targeted specificity profile were evaluated for accuracy in several ways. The referenced expression values and nucleotide contents of each considered miRNA were compared to the original data source, revealing that STREAM accurately propagates this information throughout the agent workflow and into downstream designs (Fig. 6e). Next, the relative frequency of miRNA or RBP suggestions were evaluated; in 179/180 discrete runs STREAM suggests a motif to include, with 880 total suggestions across all runs. 82% of these suggestions are miRNA binding sites, while 17.9% are RBP motifs. Only 1/880 total suggestions were not found within the available datasets (Fig. 6e). To finally assess the quality of these suggestions, a set of tissues with well-characterized, highly specific miRNAs were evaluated in more detail. These include the brain (miRNA-124)^52^, heart (miRNA-1)^53^, and colon (miRNA-192)^54^. STREAM accurately recovers these known miRNAs (Fig. 6f), while additionally identifying several supplementary candidates in both the heart and colon with high Gini coefficients relative to the target tissue and reasonably high expression values. The totality of the design pipeline averages 149.5s in runtime, compressing several months of exploratory research, component design, and final assembly. STREAM therefore provides a promising ability to systematize the hypothesis generation needed to realize the benefit of DIGs in complex settings.

## DISCUSSION

We have shown that design approaches built on top of DIGs simultaneously modulate the RL, stability, CAI, and secondary structure of an mRNA sequence under the guidance of a predictive model. By evading the requirement of differentiability, DIGs further provides a generalized way to unify diverse sources of experimental information with the benefits of model-guided sequence design while setting a new standard for runtime. These findings could be extended to other areas of biological design, including DNA and protein, while retaining a critical capability to adapt to out-of-distribution design constraints. Using this framework, forward-engineered sequences can be created without exhaustive additional screening or the computational overhead of training and fine-tuning a large model. DIGs addresses many of the hurdles that affect the deployment of machine-guided sequence design methods outside of basic research settings, offering a promising avenue to enhance their practical utility.

In designing each section of the mRNA, we uncover concepts governing the coherent assembly of de novo designed components into complete, high-performance transcripts. These findings extend our understanding of the rules guiding rational mRNA design and will be used to enhance future generative techniques and sequence-function screening experiments. Further, the results challenge the consensus regarding the role of stability on mRNA performance, a critical finding that will enhance rational mRNA design across several clinically relevant indications. The antagonistic performance of ribosomal loading– and stability-based components can now be understood through the lens of ribosomal collisions, which induces mRNA degradation^55^. Along the stability axis, collisions are avoided by recruiting a minimal number of ribosomes, which progress slowly through the highly structured UTR and CDS elements. Insertion of a high ribosome load element into a vector otherwise engineered for slow ribosomal progression leads to a mismatch between recruitment and translation rate, creating a ribosomal backlog. In contrast, high ribosomal recruitment from both UTR elements leads to strong and persistent translation if one uses a coherent, less structured coding sequence with optimal codon choices that facilitates ribosomal procession. By prioritizing interpretability, this work demonstrates how machine learning tools extend our understanding of biological principles rather than serving as a black box amplifying consensus.

Despite the promise of the DIGs technique across each of the described design tasks, some computational and biological limitations persist. In our current implementation, the objective of DIGs is to provide a single optimal sequence for the given attribute of a predictor, limiting the diversity of generated outputs. While we have shown that this approach can generalize beyond observed training samples into unexplored regions of the sequence space, creating diverse examples with similarly extreme attributes is a fundamental objective of generative design. We expect that more sophisticated programming techniques built on top of DIGs arrays will enable this task. As DIGs returns a data structure amenable with dynamic programming, it is possible that through careful algorithmic design, novel, globally optimal sequences for a given model could be returned, a guarantee that would be very hard to satisfy through other means. While coherent selection applies across many of the out-of-distribution demonstrations that we present throughout this work, we do not expect this principle to apply uniformly across the range of naturally occurring UTR sequences exhibiting both moderate ribosome load and moderate stability. The principles of orthogonal, synergistic design pathways do, however, provide a strong foundation through which a comprehensive model of UTR and CDS interaction could be constructed.

In conclusion, DIGs is an extremely efficient method for automated rational mRNA design. RNA regulatory mechanisms, out-of-distribution reporters, varying cell lines, and context-specific descriptive data is incorporated with model-based optimization through DIGs in a straightforward manner. By linking these capabilities with our agentic workflow, DIGs and STREAM greatly reduce the barrier to entry in designing high performance mRNA molecules across diverse settings and could immediately enhance many RNA therapies that are being explored in clinical applications. More generally, this work provides a tight feedback loop for converting descriptions of a biological condition of interest into potent effectors of de novo biological outputs in mRNA.

## METHODS

### SANDSTORM Datasets and Predictive Model Training

#### 5’ UTR

The SANDSTORM 5’ UTR RL model was trained as described in our previous work^4^. Briefly, the data consists of 50-nt UTR elements that were experimentally screened using polysome profiling techniques. Input data points are transformed into a paired one-hot encoding and structural contact map, at which point they are passed to the model for prediction. The model was trained for 20 epochs on this data using a train-test split of 80-20 and the Adam optimizer with a learning rate of 0.001.

#### 3’ UTR

The same SANDSTORM architecture was again applied on a previously published dataset of 3’ UTR performance^23^. In this experimental setup, both polysome profiling and mRNA abundance (used as a proxy for stability) were reported for a library of viral genomic tiles inserted into the 3’ UTR of a reporter. We reprocessed the raw read data from this study using the bowtie 2^56^ alignment program under *sensitive local* settings and filtered aligned reads for strict barcode matches using a custom python framework. For mRNA abundance, the MPRA R^57^ package was utilized to calculate RNA reads normalized to DNA reads as the target variable for model training, with normalized reads averaged across replicates. For RL, the aligned reads for each polysome fraction were again filtered for exact barcode matches and a single RL quantity was calculated using the formula from Seo et al.^23^ Input sequences were processed for prediction tasks as described above and independent model training was conducted for mRNA abundance and RL respectively using the same SANDSTORM architecture and training settings^4^.

#### CDS

SANDSTORM was utilized to estimate the MFE of CDS elements by labelling a synthetic dataset of synonymous sequences with thermodynamic models. Briefly, 500,000 random mCherry coding sequences (711 nt) or firefly Luciferase sequences (1650 nts) were created in silico by uniformly sampling from all valid codons. The secondary structure and MFE of each element was calculated using LinearFold^58^. The length-normalized MFE was calculated and carried forward as the target variable for model predictions, while the one-hot encoded sequences and structural contact maps for each CDS member were used as inputs. The model was trained with the same hyperparameter settings for 20 epochs.

#### 5’ UTR-CDS Interaction Term

Previous studies have highlighted improved RL by reducing the structural interactions of the 5’ UTR and 5’ portion of the CDS^41^. In an attempt to leverage this during DIGs design, we conducted a new training routine to predict the MFE between a specific CDS and library of synthetic UTR elements. To create labelled training data for this task, an in-silico library of 10,000 generated UTR sequences returned by the GARDN-UTR model was created spanning a range of GC content and predicted RL. We utilized NUPACK^50^ to predict the MFE between each of these UTR elements and an individual CDS element of interest, repeating this process 3 times for each of the DIGs spleen CDS elements. A SANDSTORM predictive model was then trained on the MFE value using each UTR as an input while utilizing the same training settings described previously.

### Integrated Gradients Calculations

Integrated gradients^21^ was calculated throughout this work on the one-hot-encoded sequence channel of the SANDSTORM model architectures, or on the input embeddings used in the predictive models from Zhang et al.^18^ One-hot-encoded sequence inputs transform strings of nucleotides into a vector of shape 4 × *N*, where *N* is the length of the sequence and each channel encodes a 0 or 1 to indicate the presence of a specific nucleotide at each position from 0 to *N*. Integrated gradients is defined as the path integral computed over the straight-line path between a baseline, *x*′, and a specific input sample, *x*. Assuming a differentiable predictive model *P*_δ_(*x*) accepting *i* total input dimensions, integrated gradients along the *i^th^* dimension is defined as follows:

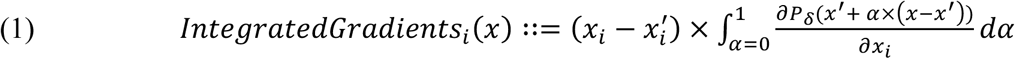

In the original integrated gradients work, *x*′ is conceptualized as a zero– or noise-vector. The resulting data structure therefore offers an explanatory map for the portions of the input, *x*, which are contributing to the output score, *P*_δ_(*x*). In this work, we reframe the technique to assign both *x* and *x*′ to be opposite and extreme examples of the feature which *P*_δ_ predicts. Without altering the computation, this shifts the interpretation of the output to an explanatory map describing the features which contribute to the opposite and extreme values of *x* and *x*′. The output of this process is therefore a static data structure encoding model preferences across the maximal observed outputs of *P*_δ_, which can be traversed as a template for designing new example data inputs with extreme features. These traversals can take the form of nearly any programming technique without the requirement of retaining differentiability. In line with this interpretation of the resulting data structure, we found that assigning *x* and *x*′ as opposite and extreme values enabled a more robust output design than assignment of *x*′ as a noise or zero vector.

For the integrated gradients calculation, we utilized between 100-250 linearly interpolated points between *x* and *x*′, creating an array of incremental intermediates between each one-hot encoded nucleotide. Additionally, in cases where the output design from our DIGs traversals returned examples with more extreme attributes than any labelled example, we repeated the process using this new example as either *x* or *x*′. We found that this process typically plateaued between 1-5 repetitions.

### Transcriptomic Analysis

Precomputed differential expression measurements were utilized in our investigations of chikungunya time course and tissue-specific expression reported by the original authors^45,46^. For time-course infections, the top N genes were ranked based on log fold change. For the tissue-specific chikungunya infection in mice and macaques, we filtered for significantly differentially expressed genes in terms of p values (p<0.05) and then utilized all upregulated and downregulated transcripts for further analysis. Endogenous CDSs for human, mouse, and macaques were exported from BIOMART^60^, with entries shorter than 100 nt pruned from further analysis. MFE values for CDS elements were calculated using LinearFold^57^. For BIOMART entries that returned multiple transcript variants, we utilized the mean MFE across reported variants while utilizing random isoform selection for codon-preference calculations. In both analyses the unexposed samples were utilized as the baseline for differential expression.

Single-cell RNA (scRNA) analysis of the mouse vaccination study^30^ was conducted using the scanpy package^61^ and the count matrix reported by the original authors. The original paper presented separate scRNA experiments from both the site of vaccination (muscle) and the lymph nodes. Differential MFE analysis was conducted using the skeletal muscle cell group following first LNP exposure as the baseline sample to isolate effects from RNA exposure. Upregulated or downregulated genes were plotted after filtering for adjusted p<0.05 and log abs(fold-change) >1. For RNA binding protein analysis, CD4+ T cells upon exposure to PBS were utilized as the baseline for differential analysis. In this case PBS was used as RNA binding proteins upregulated in response to either the LNP or mRNA would be useful for therapeutic applications. Manual processing of upregulated genes (adjusted p<0.05, log abs(fold-change) >1) was conducted to identify potential RNA binding proteins known to have a positive effect on mRNA stability or translation from the literature.

### DIGs design algorithms

#### 5’ UTR

DIGs design of 5’ UTR elements was conducted using two strategies. In the first basic example, a 50-nt 5’ UTR was created by traversing the integrated gradients array returned by the SANDSTORM 5’ UTR RL predictor in a 5’ to 3’ direction, selecting the most favored nucleotide at each position. The target and baseline samples were selected as the highest and lowest-labelled examples from the original dataset. In the second strategy, we conducted joint design based on the SANDSTORM model which predicted 5’ UTR RL and the SANDSTORM model which predicted the MFE between a library of 5’ UTRs and individual coding sequences. Using both of these DIGs arrays, we traversed from a 5’ to 3’ direction, applying a weight (alpha and beta) to the suggested nucleotide at each position from each channel, allowing us to tune the relative contributions of each in the resulting design.

#### 3’ UTR

The basic stability and RL 3’ UTRs were 170 nt in length and designed using the same 5’ to 3’ traversal that enabled 5’ UTR design, selecting the most favored nucleotide at each position. For the miRNA-142-3p UTR redesign experiments, we utilized the binding site annotations from Trobaugh et al.^27,28^ and conducted a slightly modified traversal. In this setting we populated the sequence in the 5’ to 3’ direction taking the most favored nucleotide at each position, but left the annotated binding sites unaltered. The template UTR was 353 nucleotides in length; over 2x the length of those within the training dataset used to create the DIGs template. To achieve this length in our synthetic constructs, we stitched together multiple copies of the DIGs array while retaining the binding sites in their absolute position relative to the template.

To create the synthetic CIRBP and miRNA 122 combination element, we first examined different probabilistic reconstructions of potential CIRBP binding motifs using five independent methods. First was the MEME^62^ motif automatically generated from hits listed within the CLIP-seq database entry^34^. The second was identified from Liu et al.^32^ Third was utilizing standard integrated gradients on a SANDSTORM model trained to predict CLIP-seq peaks. Fourth was using the pyhmmer^63^ toolkit to construct a profile HMM from CLIP seq binding sites. Finally, we employed a variation of the RaptGen Variational Autoencoder architecture^64^ to generate novel instances of the CIRBP binding site reconstructed from CLIP-seq data. While these methods varied greatly in their respective outputs, the motif CUCGAAGAA was consistently identified across them and used as the basis for forward design. The insertion of these elements was guided by traversing the maximum stability template from a 5’ to 3’ direction and calculating the total score implied by the insertion of the motif at each position, eventually taking the highest score. The motif was inserted into the minimum stability sequence at the corresponding location. For successive motif insertions into the UTR, the previously selected loci were omitted from future passes to prevent the inclusion of overlapping elements. We inserted two copies of the CIRBP binding site per UTR.

#### CDS

mCherry and firefly luciferase CDS designs were created using the integrated gradients array of a SANDSTORM-MFE predictive model. When designing purely with respect to structure, we traversed the array from 5’ to 3’ in incremental steps of size 3. Each possible synonymous codon for a given amino acid (*c* ∈ *C*(*a_i_*)) was indexed into this array (φ) and assigned a score based on the sum of each individual nucleotide (φ*_i,c_*). We selected the highest possible sum for inclusion into the sequence. When incorporating context-specific codon information, we supplemented this process with a custom CAI calculated from the upregulated genes identified in our transcriptomic analyses (Ψ). Each synonymous codon was now jointly considered for its impact on secondary structure by referencing the DIGs array and the favorability of the codon in the specific context through the CAI. Weight parameters were multiplied by each value to modulate the relative contribution of each information channel in the resulting design. The selection of a specific codon *c* at position *i* of the growing amino acid chain can be expressed as follows:

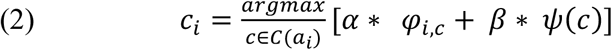

### Predictive Model Benchmarks

To compare DIGs outputs to available methods which query predictive models to inform design, we utilized several independently trained SANDSTORM models. Activation maximization, GARDN, and simulated annealing were each deployed under the supervision of a single SANDSTORM model trained to predict either 5’ UTR RL, 3’ UTR RL, or 3’ UTR stability. To assess the quality of the outputs, the predictions of each design were averaged across 10 independently trained SANDSTORM models. To create DIGs constructs, and additional 10 independent SANDSTORM models were deployed, averaging the resultant gradient maps across each before performing design by traversing through the array. In tandem, we explored simulated annealing instead using the nucleotide suggestions of the averaged DIGs array to guide sequence changes in place of the predictive model, and beam search traversals through the DIGs array to create diverse outputs. The scores of 25 examples were reported for each comparator algorithm with the time for generating this library size averaged over three repetitions.

### GEMORNA Benchmarks

To create DIGs constructs under the supervision of the predictive models published alongside GEMORNA^18^, we modified the original integrated gradients calculation to operate in embedding space. Specifically, these predictors accept tokenized embeddings of input sequences rather than raw nucleotides. We therefore computed the linear interpolation for integrated gradients directly between the embedding representations of the highest– and lowest-performing members of the original datasets (dataset X for 5′ UTRs and dataset X for 3′ UTRs). At each interpolation step, gradients with respect to the input embeddings were accumulated using the trapezoidal rule. To map the resulting attribution array back into nucleotide space, we computed the dot product between the integrated gradients vector at each sequence position and the embedding vectors corresponding to each possible nucleotide (A, G, C, U). This procedure yields a position-by-base score matrix, which we traversed greedily to generate DIGs-designed sequences. In comparison, we utilized GEMORNA specifying the ‘utr_length’ parameter as ‘long’. Output designs were scored by the predictive models published alongside GEMORNA^18^.

### In vivo Construct Design

4 unique sequences were assessed in intramuscular settings. Each sequence consisted of a synonymous Firefly Luciferase designed by different algorithmic variations alongside a set of chosen UTR elements. The first control utilized LinearDesign with a lambda value of 0 and the UTR elements from the BNT vaccine construct^65^. The second control utilized IDT’s online codon optimization protocol to create the CDS with the UTR set provided by Twist. For the experimental DIGs groups assessing muscular expression, the CAI calculated from the upregulated transcripts of genes in excised skeletal muscle tissue harvested from mice following mRNA + LNP exposure^30^ were utilized. Two distinct designs were created by stitching a CIRBP harboring UTR (Fig. 3) with a downstream RL or stability 3’ UTR. Each 3’ variant was paired with the coherent 5’ example (Fig. 4).

### STREAM Agent

The agentic workflow was implemented as a three-phase, tool-augmented pipeline to convert user prompts into tailored DIGs constructs. Implemented in python, the system supports Anthropic Claude Sonnet 4.5 and OpenAI’ ChatGPT 5.1 as LLM providers. In phase 1, an agent is provided access to the AtTract RBP database and miRNA tissue expression atlas (which contains readings for both immortalized cell lines, organs, and cell types) with several functions outlined below.

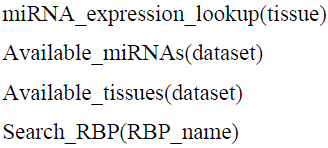

While exploring tool calls, the agent is supplemented with relevant literature via embedding-based RAG^49^ (OpenAI text-embedding-3-small, pgvector, top-k=5) with discrete chunks of every paper referenced in this work. Phase 2 of the agent creates a structured, auditable research brief that is returned to the user, maximizing exposure to the decision-making process and train-of-thought guiding the design hypothesis. Finally, phase 3 converts the research brief into a constrained recommendation, featuring 8-20 parameter recommendations for each of the 5’ UTR, CDS, and 3’ UTR. These recommendations take the form of specific parameters compatible with DIGs, including miRNA or RBP motifs to insert, stability and RL values for the UTRs, the CAI profiles presented in this work, and a tradeoff value for weighting CAI vs. structural considerations. We additionally provide a number of experimental options for altering DIGs outputs that were explored in less detail throughout this work. First, the agent can extend the length of DIGs traversals, which entails stitching together multiple copies of the precomputed arrays (similar to the approach for figure 3) to create a UTR of a desired length. We note that without proper consideration, this process can incorporate direct repeats that make synthesis and cloning difficult. Additionally, we provide options for traversing individual component arrays in a stochastic manner (Supplementary Fig. 7) with a tunable sampling temperature option to increase the diversity of outputs. Prevented patterns can similarly be specified, which deterministically block inclusion into a DIGs output by dynamically deferring to the 2^nd^ or 3^rd^ most favored nucleotide in the event that the most favored encountered during a design pass would complete the prevented pattern. Notably, exhaustive incorporation of experimental parameters into a design have not yet been fully profiled.

### Plasmid and Template Construction

All 5’ UTRs and 3’ UTRs were purchased as single stranded DNA ultramers from Integrated DNA Technologies. Coding sequences (CDS) for mCherry were purchased as gene fragments from Twist Bioscience. All 5’ UTR, 3’ UTR, and CDS combinations were cloned together with Gibson assembly (New England Biolabs) into vector backbone from Twist Biosciences. A T7 promoter sequence (GCGCTAATACGACTCACTATAGGG) was added upstream of each 5’ UTR. Plasmids were cloned in *Escherichia coli* DH5α and purified using a QIAprep Spin Miniprep Kit (Qiagen). Plasmids were linearized with PmlI and SapI for 15 h at 37 °C and purified using Monarch Spin PCR & DNA Cleanup Kit (New England Biolabs).

### mRNA Preparation

mRNA was synthesized using mMESSAGE mMACHINE T7 Transcription Kit (Thermo Fisher Scientific) with 500 ng template. IVT reactions were performed at 37°C for 2 h then treated with DNase for 15 min and post-transcriptional polyadenylation using a Poly(A) Tailing Kit (Thermo Fisher Scientific) for 45 min. mRNA was purified using Monarch Spin RNA Cleanup Kit (New England Biolabs) and eluted in RNase/DNase-free water. Quantity was determined with Nanodrop One Spectrophotometer (Thermo Fisher Scientific) and concentrations were adjusted to 500 ng/µl before storing at −80 °C.

### Cell Culture and Transfection

HEK 293T cells were grown in DMEM containing 10% FBS and 1% penicillin–streptomycin, plated at 10,000 cells per well in 96-well plates and allowed to adhere overnight. 100 ng of mRNA per well was transfected using Lipofectamine MessengerMax Transfection Reagent, according to the manufacturer’s protocol (Thermo Fisher Scientific). Six, twenty-four, and forty-eight hours later, media was removed, and cells were treated with 0.05% trypsin for 3 min to unadhere the cells from the plate. DMEM was added to stop trypsin. Cells were analyzed for mCherry expression with Attune NxT Flow Cytometer (Thermo Fisher Scientific) and analyzed using FlowJo (v10.8.1). Representative flow cytometry gating strategies can be found in Supplementary figure 8.

THP-1 cells were grown in RPMI containing 10% FBS and 1% penicillin–streptomycin in a 96-well plate at 100,000 cells per well in 200 µl media. On the same day 100 ng of mRNA was transfected per well using Lipofectamine MessengerMax Transfection Reagent, according to the manufacturer’s protocol (Thermo Fisher Scientific). Twenty-four hours later cells were analyzed for mCherry expression with Attune NxT Flow Cytometer (Thermo Fisher Scientific) and analyzed using FlowJo (v10.8.1).

Jurkat cells were grown in RPMI containing 10% FBS and 1% penicillin–streptomycin in a 96-well plate at 100,000 cells per well in 200 µl media. On the same day 100 ng of mRNA was transfected using 4D-Nucleofector X Unit (Lonza), according to the manufacturer’s protocol. Twenty-four hours later cells were analyzed for mCherry expression with Attune NxT Flow Cytometer (Thermo Fisher Scientific) and analyzed using FlowJo (v10.8.1).

### RNA and Lipid Nanoparticle Synthesis for in vivo Experiments

In vitro transcription was performed with HiScribe® T7 High Yield RNA Synthesis Kit (NEB, E2040) to synthesize mRNAs with AU cap1 analogs (Trilink, N-7114). mRNAs were polyadenylated post-IVT (Invitrogen, AM1350), and a formaldehyde denaturing gel used to confirm RNA integrity and size. RNA was encapsulated into lipid nanoparticles via rapid mixing of 1 volume aqueous RNA with 3 volumes of a lipid mixture containing SM-102, DOPE, cholesterol, and DMG-PEG2k at dissolved in ethanol at standard molar ratios of 50:10:38.5:1.5. An N:P ratio of 10:1 was used. LNP components were sterile-filtered, and synthesis performed under sterile conditions. Post-synthesis, LNPs were dialyzed against PBS overnight. RNA encapsulation was confirmed via Quantifluor assay (Promega, E3310) and LNP size measured via dynamic light scattering on a Brookhaven Nanobrook Particle Size Analyzer.

### Intramuscular Luciferase Time-Course Study

Female BALB/c mice were purchased from Jackson laboratories. Mice (n = 4) received a 50 uL IM injection into the left hindlimb muscle containing 2.5 ug of LNP-encapsulated mRNA or a PBS control. At 8-, 24-, and 72-hours post-treatment, D-luciferin (Revvity,122799) was administered IP and bioluminescence was read out 10 minutes later on an IVIS Spectrum device (PerkinElmer). ROIs were drawn to quantify photon flux for the treated muscle and mouse whole body. Due to an incomplete injection at 0 hours and complications with submandibular bleeds at 24 hours, two mice were excluded from the study.

### Code Availability

The Gardn Studio web interface and hosted agentic access point are available to academics at: https://rnadesign.app. The user interface serving complete DIGs mRNA designs is free for academic use. Agentic features are provided as a hosted service and are billed to academic users on a **per-usage basis** (to cover third-party model inference and associated compute).

## Acknowledgements

This work was supported by an NIH Director’s Transformative Research Award (R01EB037112) and startup and discretionary funds from Boston University to AAG, and by a Kilachand Award to WWW, MWG, and AAG. ATR was supported by the NIH Training Program in Quantitative Biology and Physiology (5T32GM008764). The content is solely the responsibility of the authors and does not necessarily represent the official views of the National Institutes of Health.

## REFERENCES

1. Sumi, S., Hamada, M. & Saito, H. Deep generative design of RNA family sequences. Nat Methods 21, 435–443 (2024).

2. Sample, P. J. et al. Human 5′ UTR design and variant effect prediction from a massively parallel translation assay. Nat Biotechnol 37, 803–809 (2019).

3. Wong, F. et al. Deep generative design of RNA aptamers using structural predictions. Nat Comput Sci 4, 829–839 (2024).

4. Riley, A. T., Robson, J. M., Ulanova, A. & Green, A. A. Generative and predictive neural networks for the design of functional RNA molecules. Nat Commun 16, 4155 (2025).

5. Morrow, A. K. et al. ML-driven design of 3’ UTRs for mRNA stability. 2024.10.07.616676 Preprint at 10.1101/2024.10.07.616676 (2024).

6. Iwano, N., Adachi, T., Aoki, K., Nakamura, Y. & Hamada, M. Generative aptamer discovery using RaptGen. Nat Comput Sci 2, 378–386 (2022).

7. Castillo-Hair, S. M. & Seelig, G. Machine Learning for Designing Next-Generation mRNA Therapeutics. Acc. Chem. Res. 55, 24–34 (2022).

8. Zhang, H. et al. Algorithm for optimized mRNA design improves stability and immunogenicity. Nature 621, 396–403 (2023).

9. Valeri, J. A. et al. Sequence-to-function deep learning frameworks for engineered riboregulators. Nat Commun 11, 5058 (2020).

10. Shor, J., Strand, E. & McLean, C. Y. NucleoBench: A Large-Scale Benchmark of Neural Nucleic Acid Design Algorithms. Preprint at 10.1101/2025.06.20.660785 (2025).

11. Schlusser, N., González, A., Pandey, M. & Zavolan, M. Current limitations in predicting mRNA translation with deep learning models. Genome Biology 25, 227 (2024).

12. Kukurba, K. R. & Montgomery, S. B. RNA Sequencing and Analysis. Cold Spring Harb Protoc 2015, 951–969 (2015).

13. Hong, M. et al. RNA sequencing: new technologies and applications in cancer research. Journal of Hematology & Oncology 13, 166 (2020).

14. Wang, Z., Gerstein, M. & Snyder, M. RNA-Seq: a revolutionary tool for transcriptomics. Nat Rev Genet 10, 57–63 (2009).

15. Xu, F., et al. Towards Large Reasoning Models: A Survey of Reinforced Reasoning with Large Language Models. Preprint at 10.48550/ARXIV.2501.09686 (2025).

16. Kaplan, J., et al. Scaling Laws for Neural Language Models. Preprint at 10.48550/arXiv.2001.08361 (2020).

17. Chen, J., et al. Interpretable RNA Foundation Model from Unannotated Data for Highly Accurate RNA Structure and Function Predictions. Preprint at 10.48550/arXiv.2204.00300 (2022).

18. Zhang, H. et al. Deep generative models design mRNA sequences with enhanced translational capacity and stability. Science 0, eadr8470 (2025).

19. Linder, J. & Seelig, G. Fast activation maximization for molecular sequence design. BMC Bioinformatics 22, 510 (2021).

20. Suga, K., Yamada, K. & Hamada, M. Deciphering the comprehensive relationship between 5′ UTR and 3′ UTR sequences with deep learning. 2025.05.17.654644 Preprint at 10.1101/2025.05.17.654644 (2025).

21. Leppek, K. et al. Combinatorial optimization of mRNA structure, stability, and translation for RNA-based therapeutics. Nat Commun 13, 1536 (2022).

22. Sundararajan, M., Taly, A. & Yan, Q. Axiomatic Attribution for Deep Networks. Preprint at 10.48550/arXiv.1703.01365 (2017).

23. Seo, J. J., Jung, S.-J., Yang, J., Choi, D.-E. & Kim, V. N. Functional viromic screens uncover regulatory RNA elements. Cell 186, 3291–3306.e21 (2023).

24. Barreau, C., Paillard, L. & Osborne, H. B. AU-rich elements and associated factors: are there unifying principles? Nucleic Acids Res 33, 7138–7150 (2005).

25. Tebo, J. et al. Heterogeneity in Control of mRNA Stability by AU-rich Elements *. Journal of Biological Chemistry 278, 12085–12093 (2003).

26. Kudla, G., Lipinski, L., Caffin, F., Helwak, A. & Zylicz, M. High Guanine and Cytosine Content Increases mRNA Levels in Mammalian Cells. PLOS Biology 4, e180 (2006).

27. Linder, J. et al. Interpreting Neural Networks for Biological Sequences by Learning Stochastic Masks. 2021.04.29.441979 Preprint at 10.1101/2021.04.29.441979 (2021).

28. Trobaugh, D. W. et al. Cooperativity between the 3’ untranslated region microRNA binding sites is critical for the virulence of eastern equine encephalitis virus. PLoS Pathog 15, e1007867 (2019).

29. Trobaugh, D. W. et al. RNA viruses can hijack vertebrate microRNAs to suppress innate immunity. Nature 506, 245–248 (2014).

30. Kim, S. et al. Innate immune responses against mRNA vaccine promote cellular immunity through IFN-β at the injection site. Nat Commun 15, 7226 (2024).

31. Morf, J. et al. Cold-Inducible RNA-Binding Protein Modulates Circadian Gene Expression Posttranscriptionally. Science 338, 379–383 (2012).

32. Liu, Y. et al. Cold-induced RNA-binding proteins regulate circadian gene expression by controlling alternative polyadenylation. Sci Rep 3, 2054 (2013).

33. Corre, M. & Lebreton, A. Regulation of cold-inducible RNA-binding protein (CIRBP) in response to cellular stresses. Biochimie 217, 3–9 (2024).

34. Zhao, W. et al. POSTAR3: an updated platform for exploring post-transcriptional regulation coordinated by RNA-binding proteins. Nucleic Acids Research 50, D287–D294 (2022).

35. Bicknell, A. A. et al. Attenuating ribosome load improves protein output from mRNA by limiting translation-dependent mRNA decay. Cell Rep 43, 114098 (2024).

36. Narula, A., Ellis, J., Taliaferro, J. M. & Rissland, O. S. Coding regions affect mRNA stability in human cells. RNA 25, 1751–1764 (2019).

37. Forrest, M. E. et al. Codon and amino acid content are associated with mRNA stability in mammalian cells. PLOS ONE 15, e0228730 (2020).

38. Presnyak, V. et al. Codon Optimality Is a Major Determinant of mRNA Stability. Cell 160, 1111–1124 (2015).

39. Agarwal, V. & Kelley, D. R. The genetic and biochemical determinants of mRNA degradation rates in mammals. Genome Biol 23, 245 (2022).

40. Wu, Q. et al. Translation affects mRNA stability in a codon-dependent manner in human cells. eLife 8, e45396 (2019).

41. Mauger, D. M. et al. mRNA structure regulates protein expression through changes in functional half-life. Proceedings of the National Academy of Sciences 116, 24075–24083 (2019).

42. Wu X, Bartel DP. Widespread influence of 3’-end structures on mammalian mRNA processing and stability. Cell. 2017;169:905–17.e11. 33. Van Etten J, Schagat TL, Hr

43. Zhu, Y., Zhu, L., Wang, X. & Jin, H. RNA-based therapeutics: an overview and prospectus. Cell Death Dis 13, 1–15 (2022).

44. Gray, M. et al. Chikungunya virus time course infection of human macrophages reveals intracellular signaling pathways relevant to repurposed therapeutics. PeerJ 10, e13090 (2022).

45. Bartholomeeusen, K. et al. Chikungunya fever. Nat Rev Dis Primers 9, 1–21 (2023).

46. Lin, D., et al. Multiorgan proteomic analysis of infected animal models predict potential host factors for chikungunya virus. MedComm (2020) 6, e70013 (2025).

47. Parhiz, H., Atochina-Vasserman, E. N. & Weissman, D. mRNA-based therapeutics: looking beyond COVID-19 vaccines. The Lancet 403, 1192–1204 (2024).

48. Kong, L.-Z. et al. Understanding nucleic acid sensing and its therapeutic applications. Exp Mol Med 55, 2320–2331 (2023).

49. Lewis, P., et al. Retrieval-Augmented Generation for Knowledge-Intensive NLP Tasks. Preprint at 10.48550/ARXIV.2005.11401 (2020).

50. Rishik, S., Hirsch, P., Grandke, F., Fehlmann, T. & Keller, A. miRNATissueAtlas 2025: an update to the uniformly processed and annotated human and mouse non-coding RNA tissue atlas. Nucleic Acids Research 53, D129–D137 (2025).

51. Giudice, G., Sánchez-Cabo, F., Torroja, C. & Lara-Pezzi, E. ATtRACT—a database of RNA-binding proteins and associated motifs. Database 2016, baw035 (2016).

52. Sun, Y., Luo, Z.-M., Guo, X.-M., Su, D.-F. & Liu, X. An updated role of microRNA-124 in central nervous system disorders: a review. Front. Cell. Neurosci. 9, (2015).

53. Li, J., Dong, X., Wang, Z. & Wu, J. MicroRNA-1 in Cardiac Diseases and Cancers. Korean J Physiol Pharmacol 18, 359 (2014).

54. Geng, L. et al. MicroRNA-192 suppresses liver metastasis of colon cancer. Oncogene 33, 5332–5340 (2014).

55. Simms, C. L., Yan, L. L. & Zaher, H. S. Ribosome Collision Is Critical for Quality Control during No-Go Decay. Molecular Cell 68, 361–373.e5 (2017).

56. Langmead, B. & Salzberg, S. L. Fast gapped-read alignment with Bowtie 2. Nat Methods 9, 357–359 (2012).

57. Myint, L., Avramopoulos, D. G., Goff, L. A. & Hansen, K. D. Linear models enable powerful differential activity analysis in massively parallel reporter assays. BMC Genomics 20, 209 (2019).

58. Huang, L. et al. LinearFold: linear-time approximate RNA folding by 5’-to-3’ dynamic programming and beam search. Bioinformatics 35, i295–i304 (2019).

59. Fornace, M. E. et al. NUPACK: Analysis and Design of Nucleic Acid Structures, Devices, and Systems. Preprint at 10.26434/chemrxiv-2022-xv98l (2022).

60. Smedley, D. et al. BioMart – biological queries made easy. BMC Genomics 10, 22 (2009).

61. Wolf, F. A., Angerer, P. & Theis, F. J. SCANPY: large-scale single-cell gene expression data analysis. Genome Biology 19, 15 (2018).

62. Bailey, T. L., Johnson, J., Grant, C. E. & Noble, W. S. The MEME Suite. Nucleic Acids Research 43, W39–W49 (2015).

63. Larralde, M. & Zeller, G. PyHMMER: a Python library binding to HMMER for efficient sequence analysis. Bioinformatics 39, btad214 (2023).

64. Iwano, N., Adachi, T., Aoki, K., Nakamura, Y. & Hamada, M. Generative aptamer discovery using RaptGen. Nat Comput Sci 2, 378–386 (2022).

65. Sahin, U. et al. COVID-19 vaccine BNT162b1 elicits human antibody and TH1 T cell responses. Nature 586, 594–599 (2020).

