## Supplementary Figures for "A Unified Framework for Model-Informed and Agentic RNA Design"

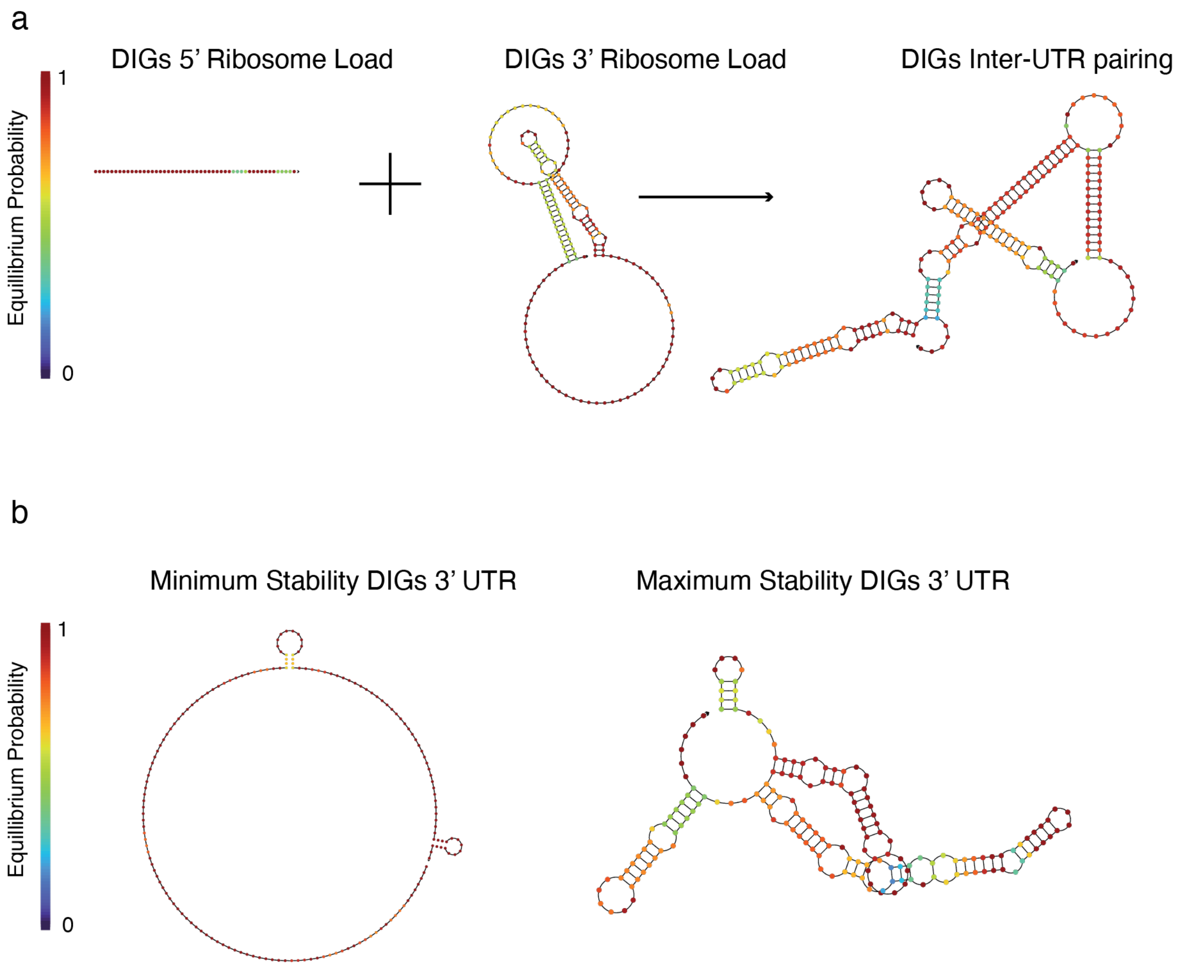


**Supplementary Figure 1 | DIGs UTRs constructs demonstrate interpretable structural features. a,** The DIGs 5' and 3' UTR elements designed to exhibit maximal ribosome loading show a strong degree of base pairing interaction, despite using independent experimental datasets to guide their generation. **b,** The maximum and minimum stability DIGs 3' UTR elements demonstrate a dramatic difference in secondary structure.

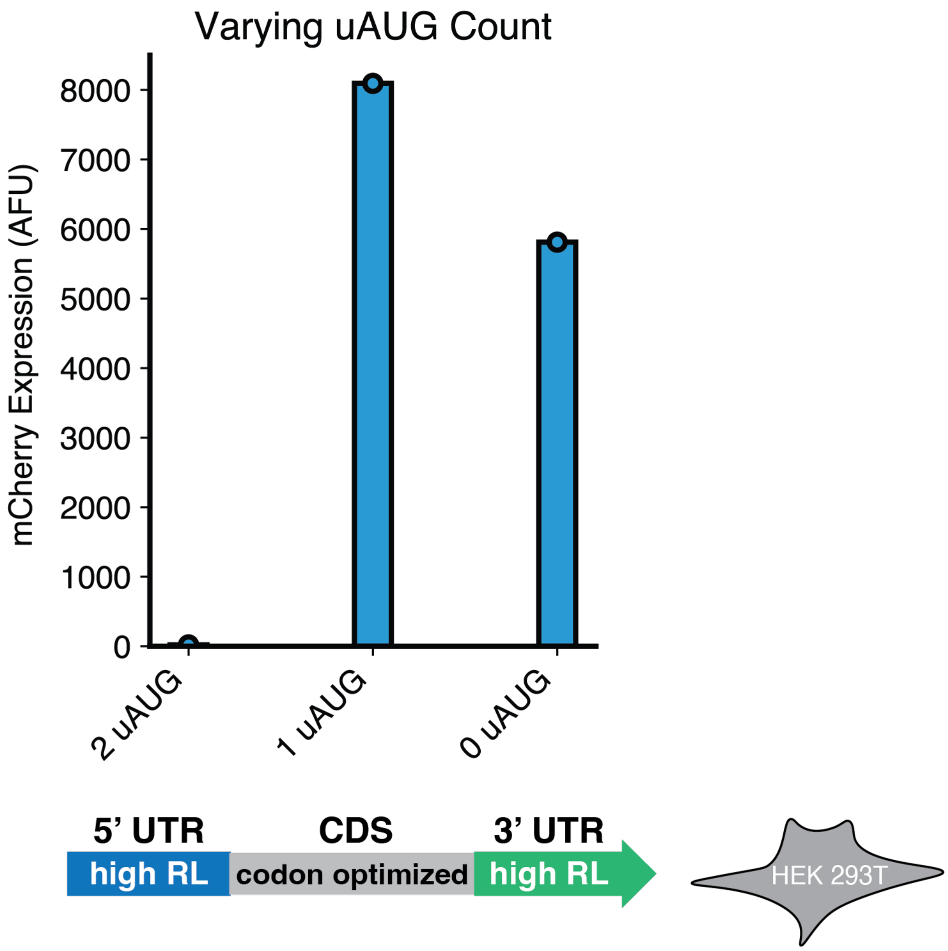


**Supplementary Figure 2 | Impact of uAUG on 5' UTR performance.** Upstream start codons (uAUGs) have been known to impact the translational capacity of mRNA molecules. By leveraging the precise grammatical control of DIGs output arrays, we created three different 5' UTR elements with similar predicted RL values that varied in their number of inserted uAUG elements. Bars denote the mean expression measurement across three biological replicates while error bars demonstrate the standard deviation.

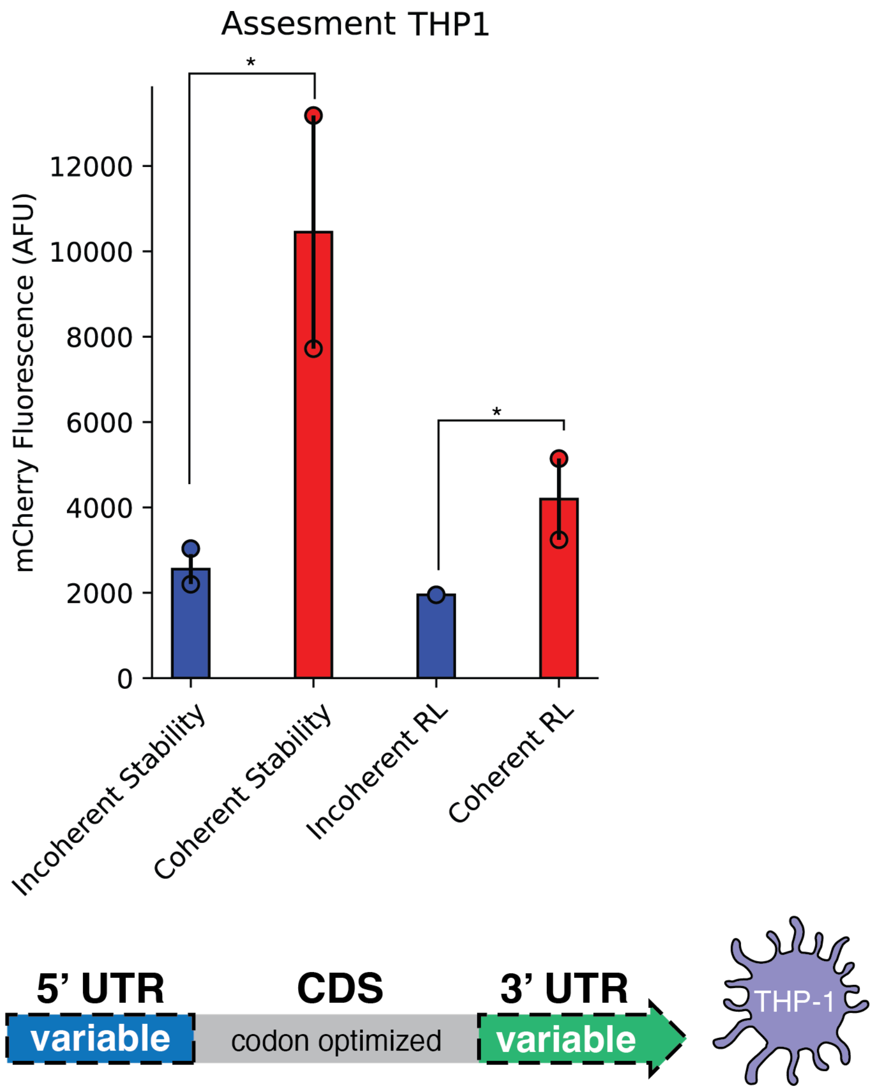


**Supplementary Figure 3 | THP-1 Coherence Assessment.** Expression measurements from a subset of the coherent and incoherent design combinations reveal that coherent selection remains informative across cell types, and also reveals that different cells may exhibit preferences for specific axes of design. Comparisons denoted ‘*’ display $p < 0.05$ measured using independent t-test for each pairwise sample comparison within the groups across three biological replicates. Data points in all plots represent the mean expression for an individual design across three biological replicates. Bars represent the mean measurements from a specific group of designs. Error bars indicate the standard deviation of the design group.

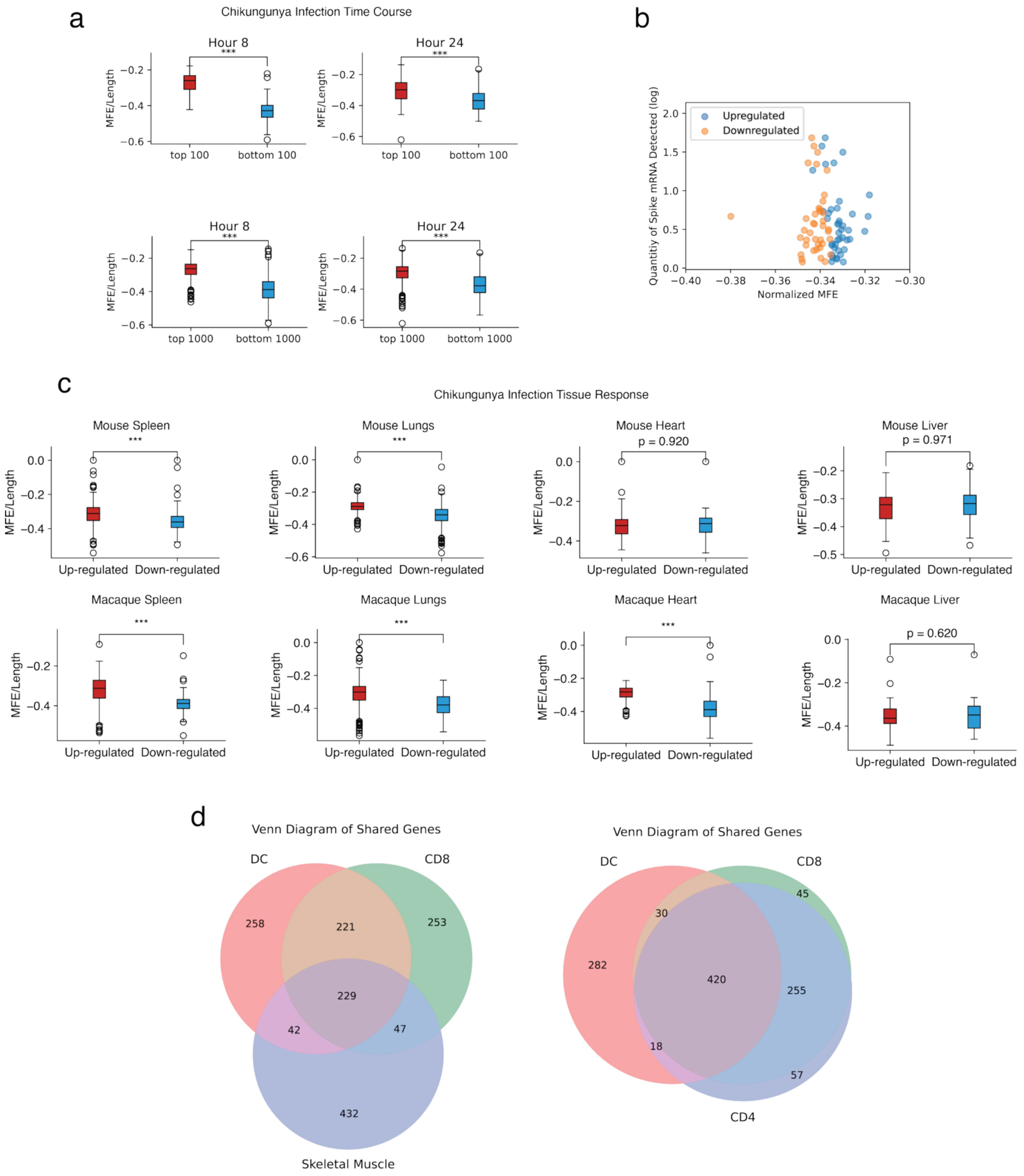


**Supplementary Figure 4 | Transcriptomic analyses uncover tissue-specific structural and codon preferences. a,** Differential expression analysis of human macrophages following chikungunya infection reveals that upregulated transcripts exhibit reduced secondary structure compared to downregulated transcripts at both 8 and 24 hours after infection. **b,** Following

COVID-19 vaccination, nearly every cell type in the murine vaccination site demonstrates a reduction in secondary structure for upregulated genes compared to down regulated genes. **c,** A tissue-specific preference for different degrees of secondary structure can be seen in both mice and macaque. In both animals, the lungs and spleen demonstrate a preference for reduced structure, while the liver demonstrates a preference for increased structure. **d,** The specific genes that are upregulated following COVID-19 vaccination show heterogeneity when comparing the muscle and spleen, motivating an exploration of tissue-specific codon preferences.

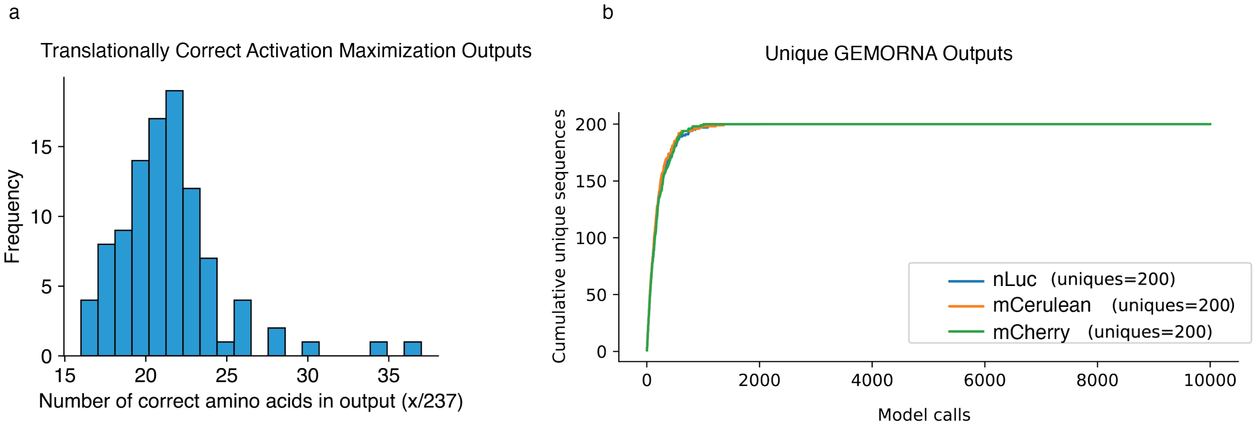


**Supplementary Figure 5 | CDS design approach assessment. a,** Activation maximization was utilized under the supervision of the predictive model for CDS secondary structure to create 100 new mCherry sequences exhibiting maximal structure. By following the gradients of this model to manipulate the structural properties of the output, most of the amino acids are changed, thereby returning a sequence that does not encode for the original protein. **b,** GEMORNA was utilized to create 10,000 examples of mCherry, 2,000 examples of nLuc, and 2,000 examples of mCerulean. The number of unique outputs was quantified, plateauing at 200 distinct examples for each target protein.

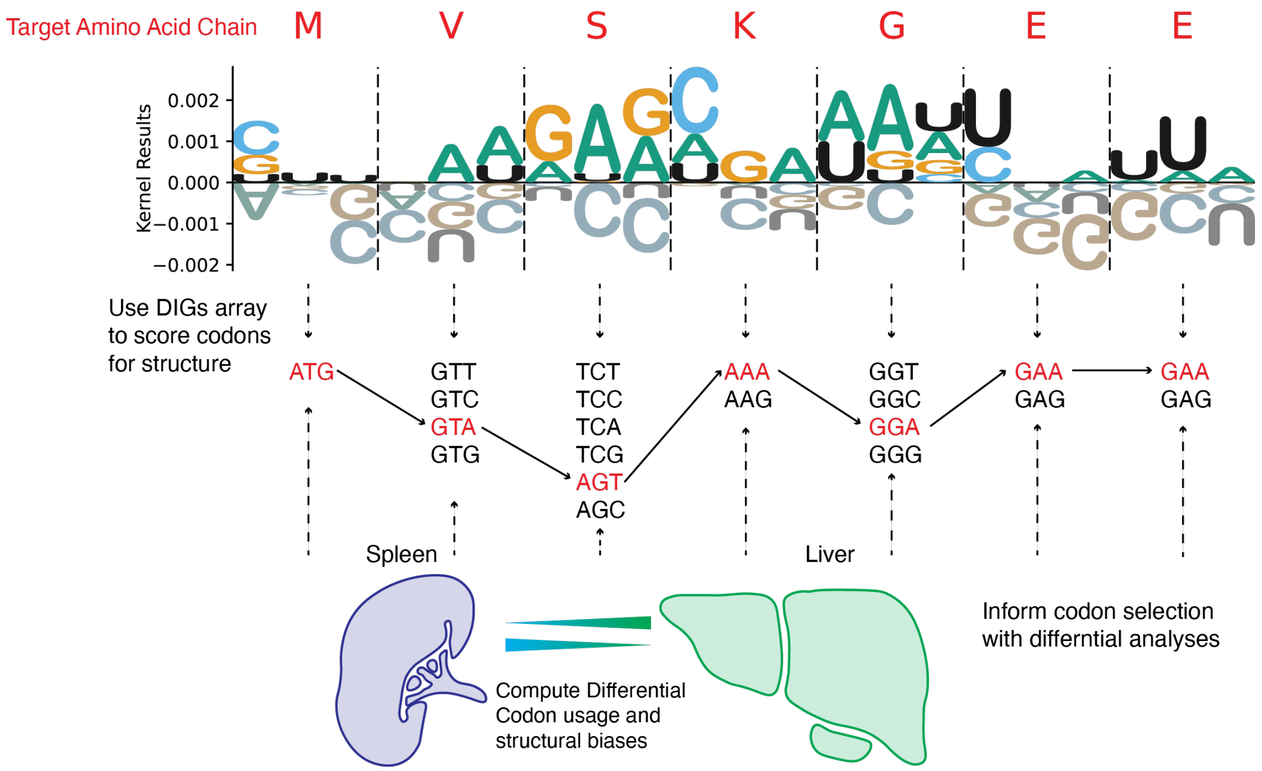


**Supplementary Figure 6 | DIGs CDS Algorithmic Implementation. a,** The DIGs array is traversed in linear time to select the optimal codon at each position based on structural considerations and the tissue-specific codon index between multiple cell types. The frozen array extracted from the predictive model is queried only for valid codons which would translate into the target amino acid. In our experiments, we deploy tissue-specific differences in codon abundance following mRNA exposure as a supplemental scoring function during design.

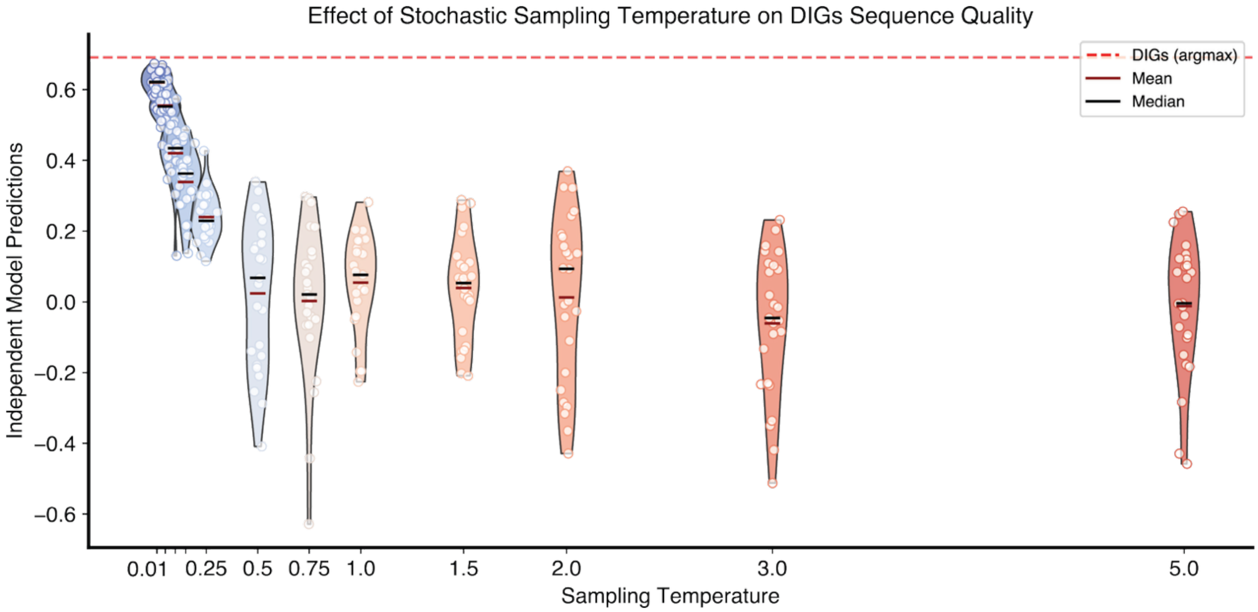


**Supplementary Figure 7 | DIGs Sampling vs. Quality.** Depicted is the effect of sampling the DIGs 3' UTR stability array with increasing stochasticity, revealing a tradeoff in sample quality and diversity. Sampling temperature is a supported experimental parameter in the online web versions of the DIGs workflow.

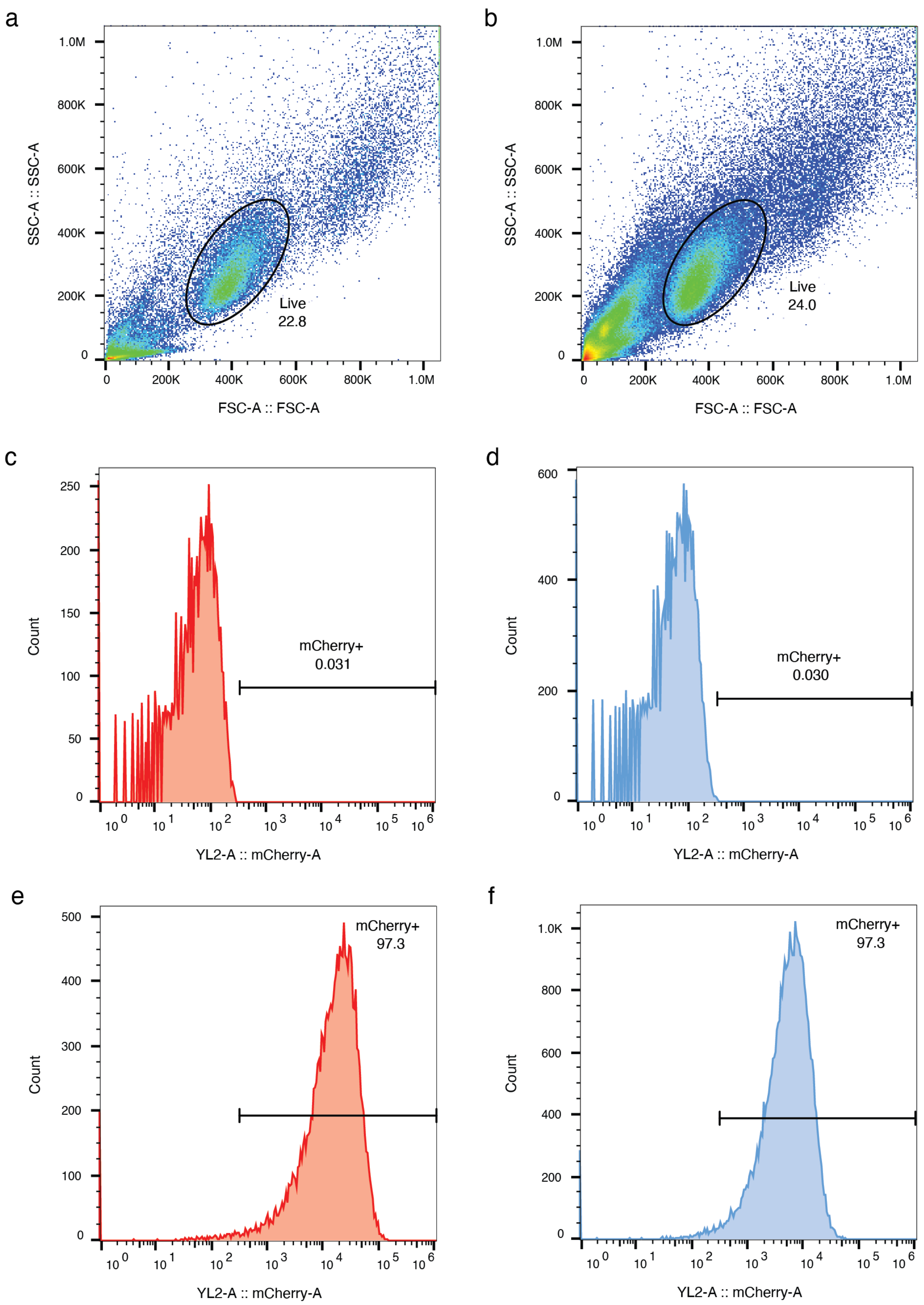
a
b
c
d
e
f
SSC-A :: SSC-A
FSC-A :: FSC-A
Live
22.8
Live
24.0
Count
YL2-A :: mCherry-A
mCherry+
0.031
mCherry+
0.030
mCherry+
97.3
mCherry+
97.3
1.0M
800K
600K
400K
200K
0
200K
400K
600K
800K
1.0M

**Supplementary Figure 8 | HEK 293T gating. a-b,** 24 hours (a) and 48 hours (b) post-transfection live gate. **c-d,** mCherry positive gate on negative control group 24 hours (c) and 48 hours (d) post-transfection. **e-f,** mCherry positive gate on OPT01 design (5' UTR: high RL; CDS: Twist mCherry codon optimized; 3' UTR: high RL) 24 hours (e) and 48 hours (f) post-transfection.
